# Integrated gene landscapes uncover multi-layered roles of repressive histone marks during mouse CNS development

**DOI:** 10.1101/2021.06.22.449386

**Authors:** Ariane Mora, Jonathan Rakar, Ignacio Monedero Cobeta, Behzad Yaghmaeian Salmani, Annika Starkenberg, Stefan Thor, Mikael Bodén

## Abstract

A prominent aspect of most, if not all, central nervous systems (CNSs) is that anterior regions (brain) are larger than posterior ones (spinal cord). Studies in *Drosophila* and mouse have revealed that the Polycomb Repressor Complex 2 (PRC2), a protein complex responsible for applying key repressive histone modifications, acts by several mechanisms to promote anterior CNS expansion. However, it is unclear what the full spectrum of PRC2 action is during embryonic CNS development and how PRC2 integrates with the epigenetic landscape. We removed PRC2 function from the developing mouse CNS, by mutating the key gene *Eed*, and generated spatio-temporal transcriptomic data. To decode the role of PRC2, we developed a method that incorporates standard statistical analyses with probabilistic deep learning to integrate the transcriptomic response to PRC2 inactivation with epigenetic information from ENCODE. This multi-variate analysis corroborates the central involvement of PRC2 in anterior CNS expansion, and reveals layered regulation via PRC2. These findings uncover a differential logic for the role of PRC2 upon functionally distinct gene categories that drive CNS anterior expansion. To support the analysis of emerging multi-modal datasets, we provide a novel bioinformatics package that integrates transcriptomic and epigenetic datasets to identify regulatory underpinnings of heterogeneous biological processes.

## Introduction

The embryonic central nervous system (CNS) is patterned along the anterior-posterior (A-P) axis, evident by the expression of brain-specific transcription factors (TFs) in anterior regions and the Hox homeotic genes in posterior regions (Figure 1A). A-P patterning of the CNS has two key consequences: first, the generation of distinct cell types in different regions, and second, the striking expansion of the brain relative to the spinal cord. Studies in *Drosophila* have revealed that anterior CNS expansion is driven by a longer phase of neural progenitor proliferation, more prevalent daughter cell divisions and faster cell cycle speeds in anterior regions, combining to generate much larger average lineages anteriorly (1). This A-P “stemness” gradient further manifests, and is driven, by an A-P gradient of neural stemness TF (e.g., SoxB family) and cell cycle gene expression (1, 2, 3). These expression gradients are in turn promoted by the selective expression of the A-P patterning TFs (3, 4, 5). However, it is unclear if the principles uncovered in *Drosophila* are fully conserved in mammals. The selective expression of TFs along the A-P axis is under control of epigenetic cues, where the Polycomb Repressive Complex 2 (PRC2) plays a prominent role (6). PRC2 mono-, di- and tri-methylates Histone 3 upon residue Lysine 27 (H3K27me1/2/3), typically resulting in proximal gene repression (Figure 1B) (7, 8).

**Fig. 1.**
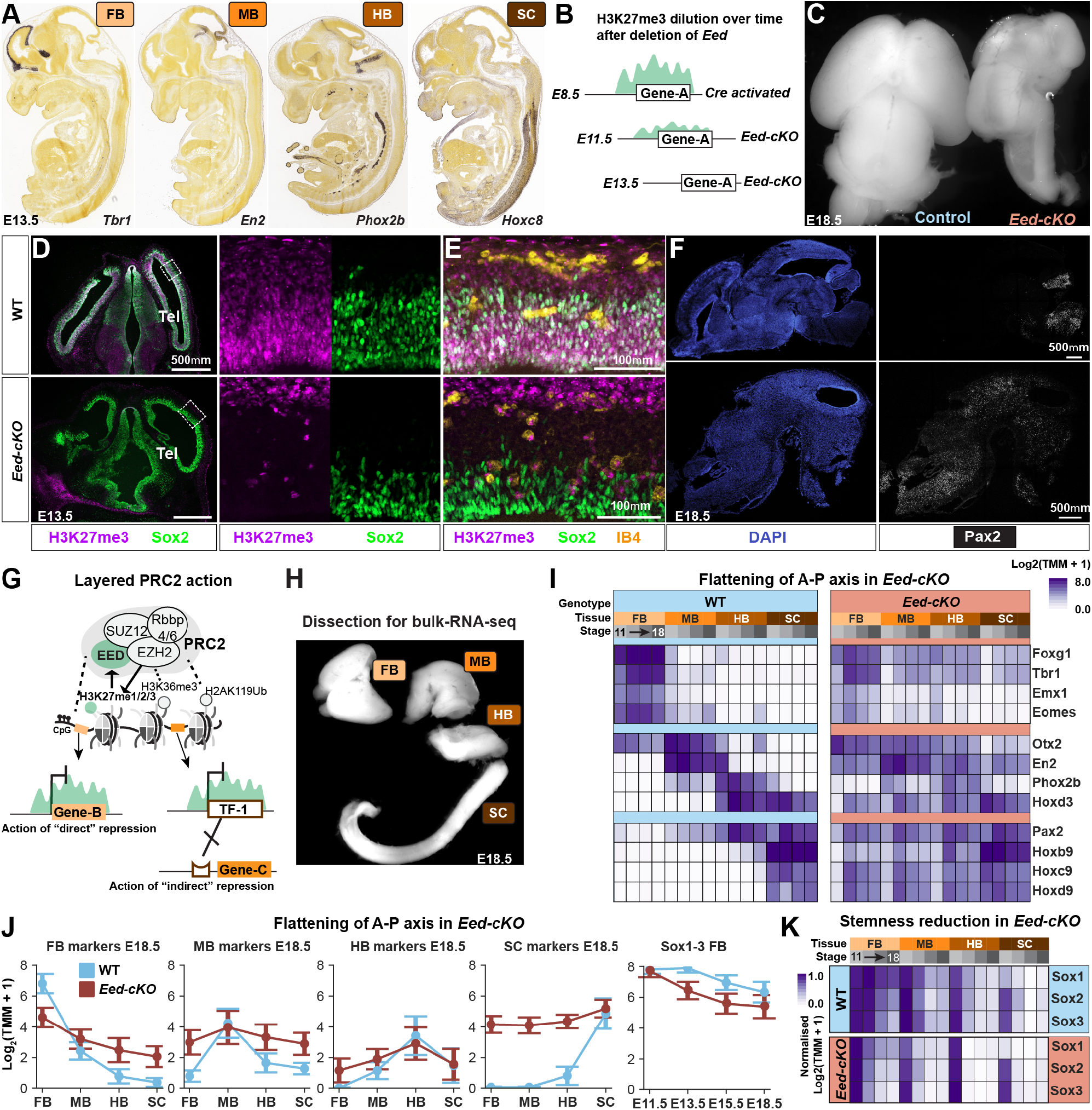
PRC2 gates the CNS A-P landscape. (A) In situ hybridization data from Allen Brain Atlas show tissue specific expression of marker genes along the AP-axis at E13.5 (B) Inactivation of PRC2, by deletion of Eed from *Sox1-Cre*, results in a gradual loss of H3K27me3. (C) WT and *Eed-cKO* littermate brains show undergrowth in the mutant. (D-E) Immunostaining for H3K27me3, Sox2 (progenitors) and IB4 (microglia) in the WT and *Eed-cKO* telencephalon at E13.5. At E13.5, deletion of *Eed^fl/fl^* by *Sox1-Cre* results in loss of H3K27me3 from the CNS and staining is only observed outside of the CNS and in infiltrating microglia. (F) Staining for DAPI (nuclei) and Pax2 in WT and *Eed-cKO* at E18.5 reveals ectopic expression of Pax2 in the entire FB and MB in the mutant. (G) The *Ezh2* methyltransferase in the PRC2 complex adds H3K27me1/2/3, which in turn interacts with the complex via EED. Other epigenetic marks are important for PRC2-chromatin interaction and/or PRC2 activity. (H) Dissection of the mouse CNS at E18.5 shows the four tissues used for bulk-RNA-seq. (I) In WT, expression of spatial marker genes is restricted to specific A-P regions, exemplified by trends in E18.5. In *Eed-cKO*, posterior genes (e.g. Hox homeotic genes) are ectopically expressed in the brain, and anterior genes are reduced in the FB and MB and ectopically expressed in the HB and SC. The effects are less pronounced at E11.5, in line with the gradual loss of H3K27me3 at E10.5-E11.5. (J) A flattening of the expression is observed (mean and standard error) for each of the marker gene groups along the A-P axis. (K) In WT, there is a spatio-temporal gradient of progenitor gene expression (*Sox1/2/3*) i.e., a gradient of “stemness” in the CNS, evident by prolonged expression of *Sox1/2/3* in the FB and MB. In *Eed-cKO*, the stemness phase in the FB and MB is shortened and the anterior CNS becomes similar to the posterior region.

Inactivating PRC2 during *Drosophila* or vertebrate CNS development, by mutating either one of the core complex genes *Ezh2* or *Eed* (*Drosophila E(z)* and *esc*, respectively), induces ectopic expression of Hox genes in the anterior CNS (5, 9), and reduces brain-specific TFs expression (4, 5). PRC2 inactivation leads to undergrowth of the anterior CNS (5, 9, 10, 11, 12), while not affecting the spinal cord growth (5). The reduced brain growth following PRC2 inactivation appears to be, at least in part, due to reduced proliferation, in particular of daughter cells (5, 13). The reduced proliferation observed in both mouse and *Drosophila* PRC2 mutants appears to result from (1) the down-regulation of brain-specific TFs, (2) upregulation of Hox genes, and (3) downregulation of neural progenitor stemness genes, such as the SoxB genes, (4) decreased expression of pro-proliferative genes and (5) increased expression of anti-proliferative genes (2, 4, 5). However, it is unclear if PRC2 acts directly and/or indirectly upon the five gene groups associated with these roles, what the full spectrum of PRC2 action is during embryonic CNS development, and how PRC2 intersects with the epigenetic landscape.

Data integration across multi-modal datasets typically occurs after statistical tests have been used to group data points, often referred to as “late integration”. However, this approach can obscure inter-dataset dependencies. In contrast, “early integration” aims to retain these dependencies, by identifying salient patterns across datasets prior to statistical analysis (14), (15). However, a lack of generalisability, interpretability and capacity to manage realistic scales of data have so far hindered widespread use of early integration across modalities (16).

Motivated by the need to integrate multiple epigenetic marks to label the chromatin landscape, ChromHMM (Chromatin Hidden Markov Model) uses a multivariate hidden Markov model. The model is trained from genome-wide assays, such as ChIP-seq of histone modifications, across conditions to capture latent chromatin states manifested in co-occurring marks (17). ChromHMM was recently used to recover two distinct states that implicate H3K27me3 during mouse embryonic development (18). However, as the transcriptome was not incorporated at an early stage it is unclear how ChromHMM chromatin states relate to gene expression during development. Moreover, how distinct chromatin states support the biological heterogeneity and the A-P patterning of the CNS was not investigated. Variational Autoencoders (VAEs) (19) are generative latent variable models able to encode relationships in mixed and multimodal data types, crossing them via successive layers of representation with “deep” learning (20, 21, 22). When “shallow” e.g., containing only a single layer, VAEs learn to map data into a lower-dimensional space akin to how principal component analysis (PCA) is commonly used. VAEs have been applied to omics data, such as single cell RNA-seq (23), bulk RNA-seq (24, 25), DNA methylation arrays (26), and histone modification ChIP-seq (27). VAEs have also been applied as an early integration method for multi-omics cancer patient data: integrating DNA methylation, bulk RNA, and copy number variation (25). However, VAEs have hitherto not been used to decode transcriptional and epigenomic events collectively, nor applied to the integration of temporal and tissue information during embryogenesis.

To understand the role of PRC2 in establishing the CNS A-P axis we generated 64 transcriptomes from wild type (WT) and PRC2 knock-out (*Eed-cKO*) mouse embryos, at different developmental stages, and from the forebrain, midbrain, hindbrain, and spinal cord regions of the CNS. We developed a workflow to analyse these data, which incorporated three stages: (1) differential analysis of transcriptomes; (2) statistical analysis of genes stratified by expression changes and wild type histone modification data; (3) VAE analysis to extract latent gene descriptors from transcriptomic and epigenetic data. The VAE analysis identified functionally heterogeneous gene cohorts with shared dependency on PRC2 and revealed the level of regulation of each gene category. Our collective findings revealed a central role of PRC2 in CNS A-P axis establishment, using a novel multi-modal, integrative approach that identifies genes driving dynamic biological processes.

## Material & Methods

### In vivo mouse models

*Eed^fl/fl^* (28) was obtained from the Jackson Laboratory Stock Center (Bar Harbor, Maine; stock number #022727). *Sox1-Cre* (29) was provided by J. Dias and J. Ericson, Karolinska Institute, Stockholm. Both lines were maintained on a B6:129 Figure S1 background. Mice were housed at the Linkoping University animal facility in accordance with regional animal ethics regulations (Dnr 69-14). Pregnant females were sacrificed and embryos dissected between stages E11.5 and E18.5. Primers used for genotyping were: Cre1: GCG GTC TGG CAG TAA AAA CTA TC. Cre2: GTG AAA CAG CAT TGC TGT CAC TT. Eed1: GGG ACG TGC TGA CAT TTT CT. Eed2: CTT GGG TGG TTT GGC TAA GA.

16 mouse embryos were extracted from 8 female mice (16 *Eed^fl/fl^*). Two embryos were extracted from each mouse at E11.5, E13.5, E15.5, and E18.5 respectively.

### RNA-seq

Mouse embryos (E11.5, E13.5, E15.5 and E18.5) were dissected to extricate the CNS (the posterior-most part of the SC was not included). The CNS was then cut into four pieces, forebrain (FB), midbrain (MB), hindbrain (HB), and spinal cord (SC) (Figure 1H). The E18.5 embryos were killed by decapitation, and then dissected (in line with ethical permits and regulations). The samples were stored at −80°C until RNA isolation, using Qiagen RNeasy Mini kit Cat.74104. RNA sequencing library preparation used the NEBNext Ultra RNA Library Prep Kit for Illumina by following manufacturer’s recommendations (NEB, Ipswich, MA, USA). The sequencing libraries were multiplexed and clustered. Samples were sequenced on Illumina HiSeq 2500, using a 50bp Single End (SE) read configuration for E13.5 embryos, 150bp Paired End (PE) read configuration for E11.5, E15.5 and E18.5, with a depth of 50-60 million reads (GeneWiz, New Jersey, NJ). The RNA-seq files are available at GEO (GSE123331). Samples from the same age were litter mates, to ensure that the WT and *Eed-cKO* are as close as possible stage-wise.

### Immunohistochemistry

Embryos were fixed for 18-36h in fresh 4% PFA at 4°*C*. After this they were transferred to 30% sucrose at 4°*C* until saturated. Embryos were embedded and frozen in OCT Tissue Tek (Sakura Finetek, Alphen aan den Rijn, Netherlands) and stored at −80°*C*. 20 and 40*μm* cryosections were captured on slides, and treated with 4% fresh PFA for 15 min at room temperature. They were thereafter blocked and processed with primary antibodies in PBS with 0.2% Triton-X100 and 4% horse serum overnight at 4°*C*. Secondary antibodies, conjugated with AMCA, FITC, Rhodamine-RedX or Cy5, were used at 1:200 (Jackson ImmunoResearch, PA, US). Slides were mounted in Vectashield (Vector, Burlingame, CA, US). Primary antibodies were: Goat *α*-Sox2 (1:250, #SC-17320, Santa Cruz Biotechnology, Santa Cruz, CA, USA), Rabbit *α*-H3K27me3 (1:500, #9733, Cell Signaling Technology, Leiden, Netherlands), Isolectin GS-IB4-ALEXA647 conjugate (“IB4”) (5-20*μg/ml*, #I32450, Molecular Probes, Thermo Fisher Scientific, Waltham, MA, USA), Rabbit anti-Pax2 (1:100, #ab232460, Abcam, Cambridge, UK). IB4 and DAPI were included in the secondary antibody solutions. Confocal microscopes (Zeiss LSM700 or Zeiss LSM800) were used for fluorescent images. Confocal series were merged using LSM software or Fiji software (30). Images and graphs were compiled in Adobe Illustrator.

### RNA-seq processing

FastQC (31) (version 0.11.9) was used to perform quality control (QC), along with multiQC (32) (version 1.8). The PE samples contained adapter content thus were trimmed using cutadapt (33) (version 2.10). Adapters used for trimming were: AGATCGGAAGAGCACACGTCTGAACTCCAGTCA (read 1) and AGATCGGAAGAGCGTCGTGTAGGGAAAGAGTGT (read 2), these were trimmed with an error tolerance of 5%, overlap of 3, and minimum Phred quality of 20. FastQC on the trimmed sequences passed QC for adapter content. RNA-seq data were then aligned to the mm10 genome using Hisat2 (34) (version 2.1.0), mm10 index was generated using the Hisat2 scripts. Reads from E13.5 were aligned using default parameters for SE reads (-U), with the other time points using default parameters for PE reads, the only parameters changed were: number of seeds set to 5; and number of primary alignments (k) also set to 5. Hisat2 reported an overall alignment rate > 90% for all files. Reads were sorted using samtools (35) (version 1.10). FeatureCounts from subread (11) was used to count the reads mapping to genes. Exon feature was used for both SE and PE reads. The PE reads were aligned such that pair fragments with both ends successfully aligned were counted without considering the fragment length constraint and excluding chimeric fragments (-p -C -B -t exon -T). Default parameters were used for the E13.5 reads (-t exon). FeatureCounts reported an average mapping to genes of ~70% for PE and ~60% for SE.

### Differential Expression

Differential expression analysis was performed using DESeq2 (36) (version 1.28.1), R (version 4.0.2). Genes were filtered if they had less than 10 counts in half of the samples. DE analysis was performed on each tissue to compare between WT and KO, where the condition (WT or *Eed-cKO*) was used as the factor and time as a batch factor. For each tissue (FB, MB, HB, SC) we used the three later time-points, E13.5, E15.5 and E18.5, for differential expression (resulting in six replicates for each test). E11.5 samples were omitted from DE owing to residual H3K27me3. We performed a similar analysis on the time points, grouping anterior tissues (FB, MB) and posterior tissues (HB, SC), resulting in eight DE analyses on WT vs *Eed-cKO* for timepoints. We performed DE between time points, such that we compare anterior time point E11.5 to anterior time point E18.5 (resulting in four replicates (two biological) for each test), using the tissue as a batch factor. Results were considered significant if a gene had an adjusted p-value of less than or equal to 0.05. Py-venn (37) (version 0.1.3) and matplotlib-venn (38) (version 0.11.5) were used for displaying Venn diagrams and seaborn (39) (version 0.10.0) was used for all other visualisations.

### ChIP-seq processing

ChIP-seq data (NarrowPeak files, IDR reproducible peaks selected) for mm10 mouse FB, MB, and HB tissues at embryonic timepoints were downloaded from ENCODE (November 2019). Peaks were annotated to entrez (40) (NCBI, database) gene IDs by using scie2g (version 1.0.0), and scibiomart (version 1.0.0) (both developed as part of the package we publish with this paper) using the annotation (mmusculus Ensembl GRCm38) from biomart, Ensembl (41). Peaks were assigned to a gene if they were located within 2.5kB upstream of the TSS or within 500bp of the gene body, except for H3K36me3 which was assigned if it fell on the gene body (upstream 2.5kB of the TSS and 500bp window after the gene ends). Peaks were retained if their adjusted p-value was less than 0.05. If multiple peaks were assigned to a gene then the peak with the greatest signal was retained. Signal and widths were recorded for each peak. If no peak was mapped to a gene, this gene was assigned a zero value. Annotations from Gorkin et al. were assigned to genes when overlapping the TSS (+-10 base pairs) using scie2g. If a gene had multiple annotations, the first one was considered, thereby reducing the number of annotated genes (by Ensembl ID) from 53,254 to 52,772. These were merged to the genes using the Entrez ID. Fisher’s Exact test in scipy (42) (version 1.5.3) was used to compare annotations between a foreground and background dataset, p-values were adjusted for multiple tests using statsmodels (43) (0.12.1) package, with alpha as 0.1 and Benjamini-Hochberg (FDR-BH) correction used.

### Label stratified analysis

Integration was performed in Python (version 3.8.2). Code and visualisations are made available and documented as a fully executable Jupyter Notebook (Jupyter Core 4.6.3). Analysis results are fully reproduced by stepping through the Notebook. Pandas (44) (version 1.0.3) was used to merge the FeatureCounts files on Entrez gene ID, yielding a dataset of 27,179 rows. Gene names were annotated to merged data frame using ensembl mappings from entrez to gene name, from this, there were 6279 genes without gene names (predicted or nc-RNA), which were omitted from the subsequent analysis, leaving 20900 genes. RNA-seq data were normalised by using EdgeR’s (45) (version 3.30.3) TMM method, the log2 + 1 was then taken of the TMM counts using numpy (46) (version 1.18.2). Peak data were merged on assigned entrez ID as per the ChIP-processing section above.

We performed a simple stratification to annotate genes based on changes in expression and repressive mark presence. We labelled each gene as unaffected, partly affected or consistently affected, by using the expression response to PRC2 in-activation as per the DE analyses. Partly affected genes refers to genes displaying a significant difference in expression, an absolute log_2_ *FC* greater than 1.0, between WT and *Eed-cKO* in one to three of the DE analyses of FB, MB, HB, SC, or the anterior temporal analyses (E11.5, E13.5, E15.5, E18.5). Consistently affected refers to genes exhibiting an absolute log_2_ *FC* > 1.0 in > 3 of the eight DE analyses. To annotate genes with WT histone modification profiles we collected publicly available WT ChIP-seq H3K27me3 data for a range of embryonic mouse tissues (47). We then labelled each gene as marked or unmarked based on the presence of a H3K27me3 peak in WT, E16.5 ChIP-seq within 2.5kB of the transcriptional start site (TSS).

### VAE analysis

VAEs are implemented in scivae (developed by us for this project) (version 1.0.0), which in turn uses tensorflow (48) (version 2.3.1) and Keras (49) (version 2.4.3). VAEs were created using the consistently affected genes as input (randomly sub-divided into a training set with 85% genes). Input was the normalised transcriptome (64 features), the log2 of the H3K27me3 signal (21 features), and the log fold change from the DE analyses was used (12 features). All input data were normalised between 0 and 1. Mean squared error was used as the loss metric, with MMD (kernel) as the distance metric for the sampling function, with a weight of 1.0 was used. The VAE was trained for 250 epochs using a batch size of 50. Different numbers of latent nodes were tested, ranging from 1 to 32. Selu activation functions were used for the first input and final output layers with Relu used for internal layers; adam optimiser was used with parameters: beta1 = 0.9, beta2 = 0.999, decay = 0.01, and a learning rate of 0.01. Gene cohorts were calculated for each latent dimension from the 3 node, consistently affected dataset, with genes having a value greater than one standard deviation (+1.25SD) from the mean (0). This resulted in six gene cohorts (two for each node) with 152, 238, 154, 352, 214 and 187 genes, respectively, these were used in subsequent functional analyses. For comparisons to other methods, PCA and tSNE from sklearn (version 0.0) was used, UMAP from umap-learn (50) (version 0.4.2), PHATE from phate (51) (version 1.0.7). Default parameters were used, except for changing the number of components (3 or 6) and for tSNE, running “method=exact”, when n_components=6. Given tSNE, UMAP and the VAEs may vary in terms of projection based on the initiating seed 30 runs were completed. For each of these tools a random seed was generated per iteration and then the separability was quantified. Separability was determined as able to put gene markers from the same tissue “near” and those from less similar tissues “far away”. Far away is defined as having a significant difference between the within cluster distance (sum of square differences to the mean) and between cluster distance (pooling the two groups). The % correct out of the 30 runs were reported for each tool. For PCA (deterministic) and PHATE (while PHATE did allow for a seed to be set the result was the same for each iteration) a binary result was reported for each separability metric.

### Functional analysis

Over representation analysis on the gene cohorts was performed in R using enrichGO from clusterprofiler (52), (version 3.16.1). Entrez IDs were used and BH correction with FDR alpha of 0.1, using all GO annotations. Gene set enrichment analysis was performed using fgsea (53) (version 1.14.0), genes were ranked by each of the VAEs latent dimensions.

### Reproducible, generative methods applicable for other dynamic systems

Our model of the developing mouse CNS is available as a downloadable package where the profile of any mouse gene can be queried in terms of its PRC2 response. In addition to this, we provide a Python package with tutorials in R and Python for using the VAE for other dynamic systems where researchers are interested in integrating epigenetic and expression information. Our packages have been optimised for reproducibility by enabling saving of the VAE state, visualisation, and logging.

## Results

### PRC2 is critical for the anterior-posterior CNS axis

To inactivate the PRC2 complex in the mouse CNS, *Sox1-Cre* was used to conditionally delete *Eed^fl/fl^* in the CNS (denoted *Eed-cKO* herein). This resulted in the inactivation of *Eed* at E8.5, with a gradual reduction of the H3K27me3 mark (Figure 1B), presumably due to replication-mediated dilution, until it is undetectable by immunostaining in the CNS proper, at E11.5 (5), visualized at E13.5 herein (Figure 1D-F). *Eed-cKO* embryos displayed a striking up-regulation of posterior genes, such as *Pax2* (Figure 1F), in the FB/MB, and severe brain underdevelopment (Figure 1C), in large part due to a truncated proliferation phase (5).

We conducted a total of 64 bulk-RNA-seq experiments across wild type (*Eed^fl/fl^*; referred to as WT) and *Eed-cKO*, at four developmental stages; E11.5, E13.5, E15.5 and E18.5, and of four tissues: forebrain (FB), midbrain (MB), hindbrain (HB), and spinal cord (SC) (Figure 1G). Analysing the WT bulk-RNA-seq data for expression of the neural progenitor stemness genes *Sox1/2/3* (54) underscored the spatiotemporal stemness gradient (Figure 1I-J). In *Eed-cKO*, both the FB and MB displayed a more rapid downregulation of *Sox1/2/3*, while the HB and SC were less affected (Figure 1I). Analysis of spatial TF markers in WT revealed the expected selective gene expression (Figure 1A) along the A-P axis (Figure 1H). In contrast, in *Eed-cKO* mutants FB, MB and HB markers were downregulated in their specific regions, and ectopically upregulated in adjacent regions (Figure 1H-I). SC markers (e.g., Hox genes) were ectopically expressed in all anterior regions (Figure 1H-I). The mutant effects were less pronounced at E11.5 (Figure 1H), in line with the gradual loss of the H3K27me3 mark during E10.5-E11.5 (5). These results revealed that *Eed-cKO* mutants displayed a striking “flattening” of the CNS A-P axis, evident from the downregulation of brain TFs, the ectopic expression of Hox genes in the brain, and anterior downregulation of stemness genes.

### PRC2 inactivation results in posteriorization of the anterior CNS

Analysing the global gene expression differences along the CNS A-P axis, we found major differences in the baseline WT transcriptomes, with FB and MB being strikingly different from the SC (Figure 2A). When compared to SC, the FB showed 4,771 differentially expressed genes (DEGs) and MB 2,555 DEGs (log_2_ *FC* > 0.5, *P* < 0.05, where log_2_ *FC* is the log2 transformed fold change; pooled time-points) (Figure 2A). In addition, all other comparisons revealed substantial gene expression differences, underscoring the uniqueness of each axial level (Figure 2A).

**Fig. 2.**
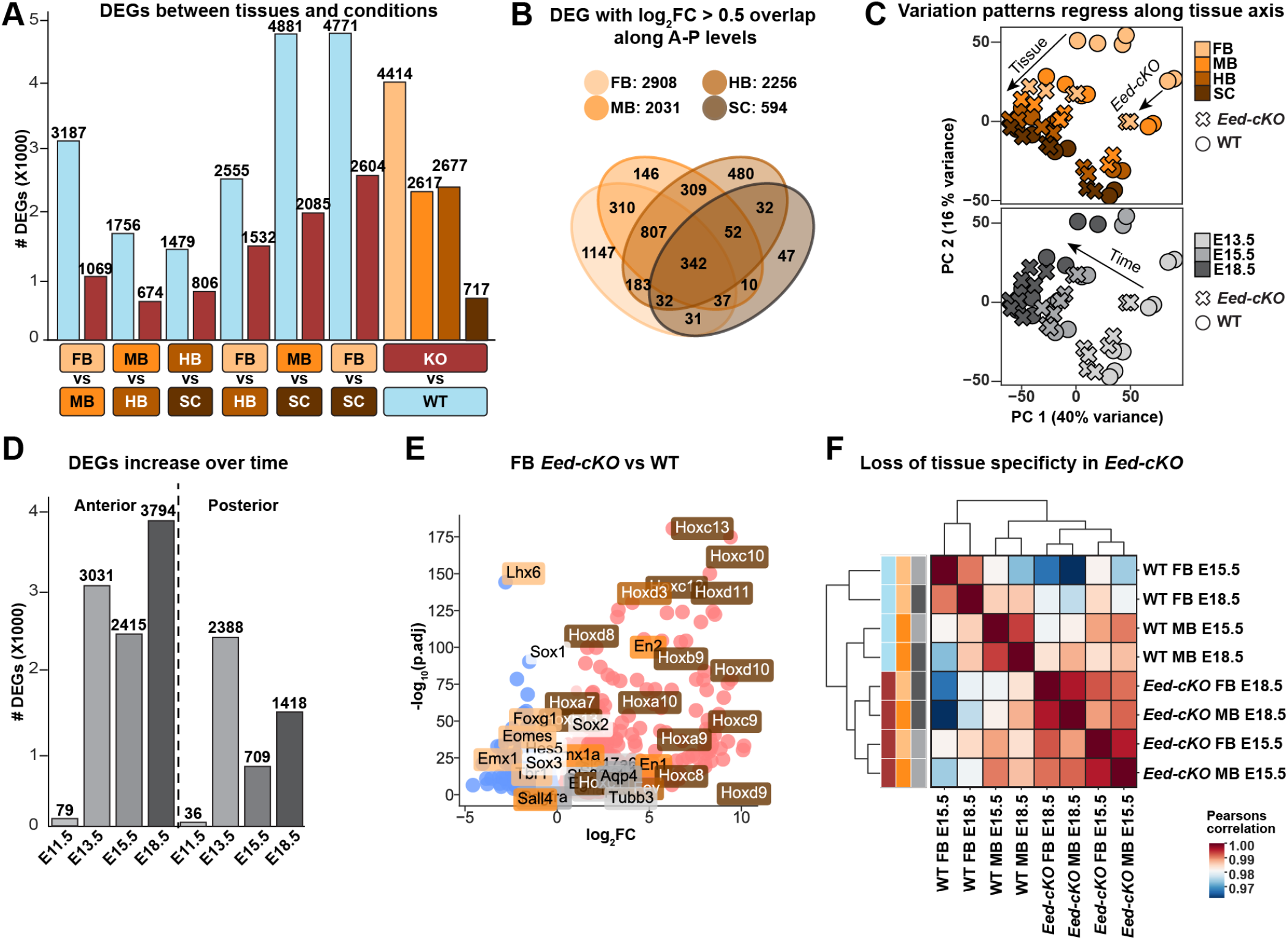
*Eed-cKO* mutation flattens the CNS A-P gradient. (A; left side) Comparative DEG analysis between different CNS levels, in WT and *Eed-cKO*, based upon pooled time points. In WT, adjacent CNS tissues display fewer differences than distal ones, especially when compared to SC (DEGs; log_2_ *FC* > 0.5, *P* < 0.05, where log_2_ *FC* is the log2 transformed change between mutant and WT). In *Eed-cKO*, the number of comparative DEGs are strongly reduced. (right side) *Eed-cKO* strongly affects anterior tissues e.g., FB (4,414 DEGs) while the SC is less affected (717 DEGs). (B) Venn diagram depicting DEG overlap in WT vs. *Eed-cKO* for each tissue. FB is most severely affected while more posterior tissues display more overlap in their response. (C) PCA of normalised RNA-seq count profiles labelled with tissue, condition (WT vs *Eed-cKO*) and time (E13.5–E18.5; E11.5 omitted); arrows indicate the A-P shift induced by Eed mutation. (D) DEG analysis over time for combined anterior (FB and MB) and posterior (HB and SC) tissues reveals that *Eed-cKO* affects the anterior more than the posterior CNS, and that the effects increase over time. (E) Volcano plot of *Eed-cKO* vs. WT expression in FB, showing that Hox genes are upregulated while brain-specific genes are downregulated. (F) Correlation between late stage anterior tissues showing a reduction of tissue specificity in *Eed-cKO*.

These axial differences were reduced in *Eed-cKO* mutants, with gene expression differences almost halved when comparing FB to SC, and MB to SC (Figure 2A). The FB was most affected, with 4,414 DEGs, while SC displayed considerably smaller effects, with only 717 DEGs (Figures 2A, S1, S2, S3). Surprisingly, only 342 genes were shared across all four tissue analyses, indicating that for the majority of DEGs the role of PRC2 is specific to each axial level (Figure 2B). While PRC2 inactivation generally caused upregulation (e.g., Hox genes), analysis of the FB revealed that a number of brain-specific TFs were downregulated (Figure 2E).

To investigate temporal variation throughout development, we grouped the FB and MB into “anterior”, and the HB and SC into “posterior” sections. This revealed that the A-P axis differences between WT and *Eed-cKO* were most pronounced in the anterior CNS at E18.5 (Figure 2D). However, the largest increase in DEGs occurred between E11.5 and E13.5, in both the anterior and posterior tissues (Figure 2D).

In line with the effects on specific marker genes (Figure 1H-J), PCA (Figure 2C) revealed transcriptome wide changes supporting the notion that *Eed-cKO* mutants were posteriorized along the A-P axis. Specifically, the mutant FB transcriptome was more similar to the WT MB, and the mutant MB to the WT HB. This trend was most evident at E13.5 but observed at all later stages (Figure 2C). The posteriorization of each *Eed-cKO* tissue was also evidenced by quantification of the normalised sum of square differences between the transcriptomes (Table S2) and correlation between anterior samples (Figure 2F).

### PRC2 inactivation does not trigger extensive ectopic expression of non-CNS genes

To address if CNS-specific PRC2 inactivation resulted in ectopic expression of peripherally expressed genes, we surveyed for genes that were not expressed in the CNS at any axial level or stage but were activated in *Eed-cKO* mutants. Somewhat surprisingly, we only identified 213 genes in this category (Figure S4, Table S2). Hence, in contrast to the extensive A-P gene expression changes within the CNS, inactivation of PRC2 in the CNS did not result in widespread breakdown of germ layer barriers of gene expression (Figure S4).

### H3K27me3 only partly explains widespread effects of PRC2 inactivation

Publicly available WT H3K27me3 ChIP-seq data (47) revealed that the region- and gene-specific profiles for several of the spatially restricted genes, *Foxg1*, *En2* and *Hoxc9*, were consistent with a direct repressive role of the PRC2 complex (Figure 3A). However, gene expression changes caused by PRC2 inactivation may result from layers of regulation when considered across the developmental trajectory. To begin addressing this issue in a systematic manner, we performed a “label-stratified” analysis, comparing gene expression with histone modification profiles. We labelled each gene as unaffected, partly affected or consistently affected, by using the expression response to PRC2 inactivation as per the DE analyses (Figure 3B). Thereby, the H3K27me3 state and expression response to *Eed-cKO* jointly defined six exclusive categories of genes (Figure 3B), see Methods for details.

**Fig. 3.**
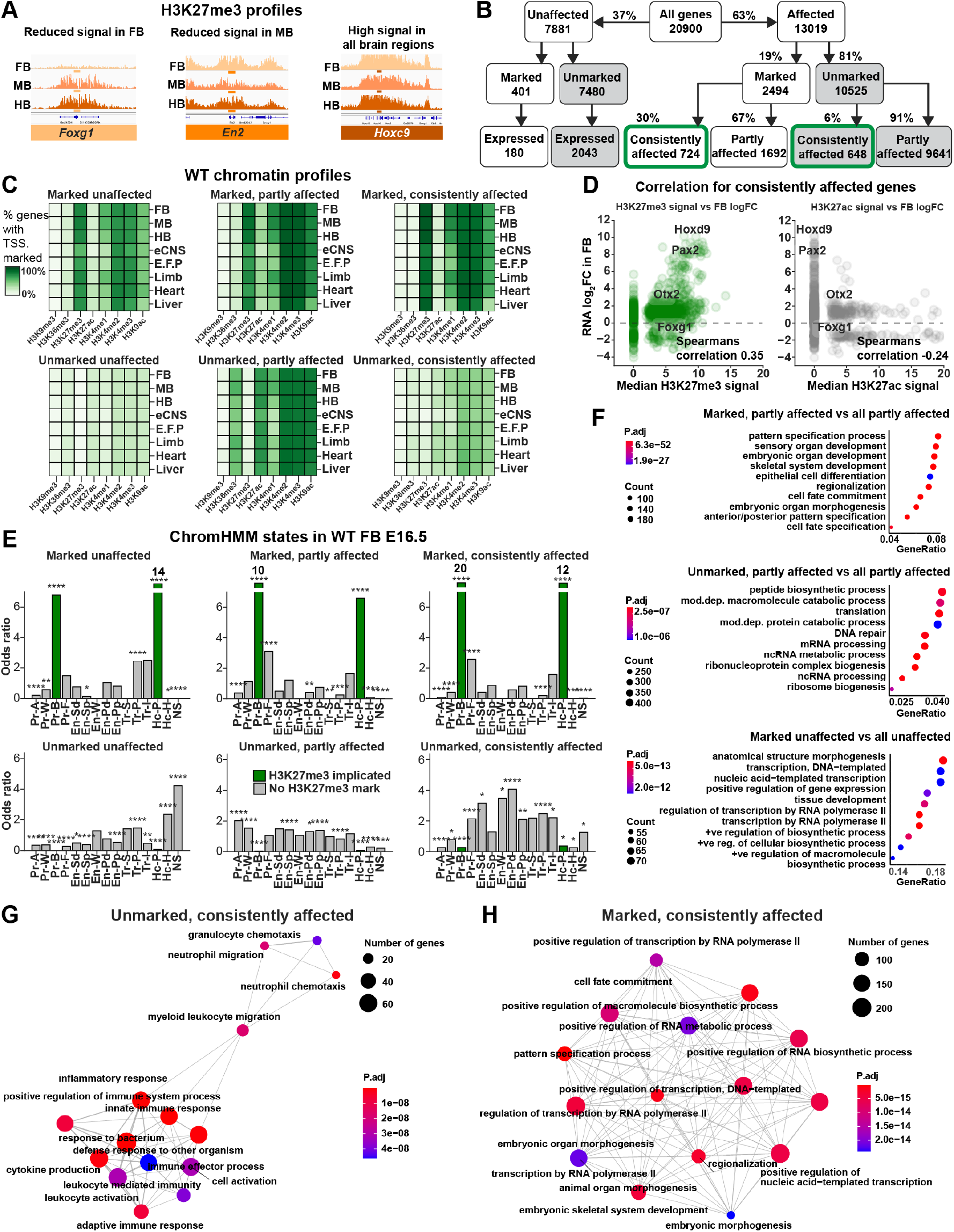
Direct and indirect control by PRC2. (A) H3K27me3 signal for three markers genes, at E16.5 in WT, showing correlation between H3K27me3 and expression patterns. (B) Gene categorising flow based upon the expression response to *Eed-cKO* mutation and the presence of H3K27me3. (C) Heatmaps showing the percentage of genes with different histone modifications at the TSS for each defined gene category, in the CNS and reference tissues. (D) Correlation between FB log_2_ *FC* and repressive (H3K27me3) and activating (H3K27ac) marks, of the consistently affected gene category, reveals limited correlation. (E) Enrichment of ChromHMM chromatin states, expressed as the odds ratio between positives in each gene category vs. all genes. (F) Functional Gene Ontology (GO) enrichment analysis by clusterProfiler of “partly affected” and “unaffected” genes shows greater similarity between “marked” groups irrespective of repsonse to PRC2. (G) GO enrichment of “Consistently affected” and “unmarked” genes reveals “inflammatory response” and “positive regulation of immune system process” (edges represent shared genes between GO terms). (H) GO enrichment analysis of “consistently affected” and “marked” genes identifies terms related to embryonic development along the A-P axis.

Given PRC2’s role in maintaining tissue specificity, we hypothesised that genes in each category would display chromatin profiles that were specific to tissue. Within each gene category histone modifications in FB, MB, HB, neural crest (eCNS), embryonic facial prominence (E.F.P), limb, heart and liver, at E16.5 (47) were surprisingly similar, but between categories differences appeared (Figure 3C). Specifically, genes in the three H3K27me3-marked categories were commonly marked with H3K4me2/3 marks, but not H3K36me3, indicating their bivalent status (Figure 3C). Within the three H3K27me3-unmarked categories, active marks (H3K36me3, H3K27ac, H3K4me2/3) were primarily observed in the partly affected category (Figure 3C).

To further investigate the relationship between the H3K27me3 mark and gene expression, we analysed the correlation between FB log_2_ *FC* and H3K27me3 signal in the “consistently affected” gene category (Figure 3D). We found a limited positive correlation (*ρ* = 0.35, *P* < 0.01) between H3K27me3 and FB log_2_ *FC*. This exceeded the correlation between the FB log_2_ *FC* and H3K27ac signal, which reported minimal negative correlation (*ρ* = −0.24, *P* < 0.01).

The limited correlation between H3K27me3 and gene expression prompted us to investigate whether a combination of histone marks i.e., chromatin states in a gene promoter could provide greater insight into the FB gene response to PRC2 inactivation. To this end, we assigned ChromHMM-predicted epigenetic states to each gene, based upon FB at E16.5 (18). We found that genes in all three H3K27me3-marked categories (unaffected, partly affected and consistently affected) exhibited similar epigenetic states, with strong signals for the Pr-B (Promoter-Bivalent) and Hc-P (Heterochromatin-Permissive) states (Figure 3E). The enrichment of these H3K27me3-implicated states was considerably higher for the consistently affected category, showing that if a gene is marked by H3K27me3 at E16.5 in the FB, it is likely to be affected by knocking out Eed (Figure 3E). The consistently affected unmarked genes stood out, with higher enrichment of enhancer states: En-Sd (Enhancer-Strong-TSS-distal), En-W (Enhancer-Weak-TSS-distal) and En-PD (Enhancer-Poised-TSS-distal) (Figure 3E), suggesting that these genes are indirectly regulated by PRC2. There were approximately equivalent numbers of genes in the consistently affected categories that were unmarked and marked, indicating that lack of the H3K27me3 mark did not rule out effects in *Eed-cKO*.

To understand the functional heterogeneity of genes within and between each category, we also tested for overrepresented Gene Ontology (GO) terms associated with their proteins (Figure 3F). For the H3K27me3-marked, both partly and consistently affected genes were enriched for A-P axis related terms, (e.g., pattern specification). Surprisingly, H3K27me3-marked but unaffected genes were also enriched for regulation and development, indicating that not all marked developmental genes were affected by *Eed-cKO* (Figure 3F). Unmarked and partly affected genes were associated with RNA processing terms (Figure 3F), while the unmarked, consistently affected genes were primarily enriched for immune response genes (Figure 3G-H).

### Variational Autoencoder finds latent codes for mixture of features

Genes marked by H3K27me3 in the CNS tended to be affected by *Eed-cKO*. However, the effect on unmarked genes was nebulous e.g., correlating the log_2_ *FC* of the response with the experiment-wide median H3K27me3 state revealed only a weak correlation (*ρ* = 0.35, *P* < 0.01), underscoring the limited ability of the H3K27me3 mark alone for predicting the expression response to *Eed-cKO* (4A). These findings suggested that labelling genes without jointly including details of developmental stage and tissue obscured features required to identify co-regulated genes.

To more comprehensively understand gene co-regulation, while avoiding exhaustively screening all possible permutations of features, we developed an approach using a variational autoencoder (VAE). Similar to PCA, Uniform Manifold Approximation and Projection (UMAP) (50), t-Distributed Stochastic Neighbour Embedding (tSNE) (55), and Potential of Heat-diffusion for Affinity-based Transition Embedding (PHATE) (51), VAEs map data to a lower-dimensional latent space to thereby facilitate interpretation (Figure S5). We opted to use a VAE to integrate the signal and response data owing to VAEs reported abilities of extracting biologically meaningful features from non-linearly dependent data (24). We defined a PRC2 profile of prioritised features (97) representing each gene for input to the VAE: RNA-seq data for WT and *Eed-cKO*, and WT H3K27me3 signal. Our goal was to integrate the data into a relatively small set of features and use the model to interrogate relationships between the WT H3K27me3 signal and the gene expression response to *Eed-cKO* (Figure 4B).

**Fig. 4.**
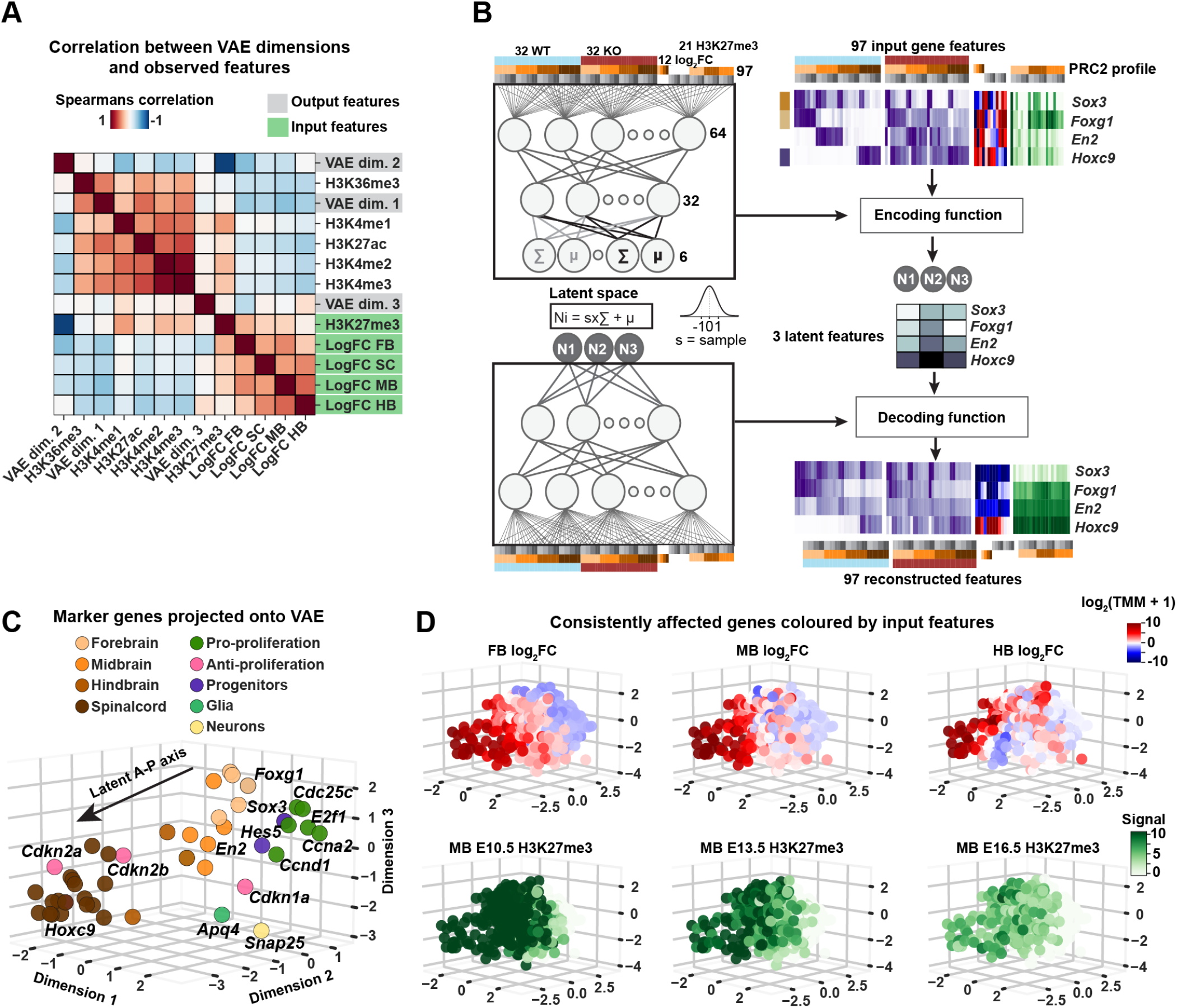
Integrated PRC2 profile forms latent A-P axis. (A) Heatmap of Spearman correlation coefficients across all consistently affected genes between VAE latent codes and selected annotations. (B) Simplified VAE model and the gene-specific input to the VAE, supplying a range of experimental observations (example genes shown) to a (non-linear, trainable) “encoding function”, which defines a latent code for each gene. A “decoding function” is trained to reconstruct profiles for each gene, subject to VAE constraints. (C) Selected marker genes plotted in VAE D=3 latent space, showing an A-P gradient. (D) Consistently affected genes plotted in VAE latent space, coloured (top row) by log_2_ *FC Eed-cKO* vs. WT in FB, MB, and HB, and (bottom row) the median signal in H3K27me3 across development.

VAE was trained on PRC2 profiles, to find a latent “code” for each gene deemed salient once all data points were considered (Figure 4B). Because the results were reproducible and robust to parameter perturbations, the VAE architecture and parameters were chosen with minimal tuning. When the VAE used three or more hidden nodes, referred to as latent dimensions (D), we observed only minor reconstruction loss (Figure S4G). Hence, at *D* ≥ 3, intermediate layers captured sufficient information to successfully decode essential variation across the full data set. We tested two versions of the data set: the 1,159 consistently affected genes (as defined above), and all of the 13,019 affected genes, with D=3 and D=6. We found that the consistently affected genes provided the better training data set; data points occupying the “middle ground” had little effect on the resulting organisation of statistically highlighted genes, but challenged ocular assessment (Table S2, Figure S5A-B). Based upon these findings, we subsequently used the VAE with *D* = 3, trained with the consistently affected gene data set.

### VAE latent code places genes along A-P axis

While no pair of VAE dimensions correlated measurably (|*ρ*| < 0.1 for all pairs), as anticipated, each VAE dimension correlated with a number of input features (Figure 4A). For instance, dimension 2 correlated negatively with the median H3K27me3 signal (*ρ* = −0.93, *P* < 0.01) and weakly with FB log_2_ *FC* (*ρ* = −0.42, *P* < 0.01). This basic trend was easier to access in the model compared with FB log_2_ *FC* and median H3K27me3 when measured directly and linearly (*ρ* = 0.35, *P* < 0.01).

To further validate that the VAE latent code uncovered biologically relevant CNS features, we tracked the aforementioned marker genes (Table S1). We measured their separability as distinct groups, and noted that the FB, MB, HB and SC genes were placed along a latent version of the A-P axis (Figure 4C). In addition, investigating the placement of proliferation genes (Table S1) we noticed that pro-proliferative genes were placed in the vicinity of FB genes, while antiproliferative genes were placed adjacent to the SC genes, in agreement with the enhanced anterior proliferation (Figure 4C). Logically, markers for neurons and glia did not group along the A-P axis, in line with the generation of these cell types at all axial levels during the embryonic stages analyzed (Figure 4C). We confirmed VAE codes at *D* = 6, finding that both configurations distinguished between the chosen marker gene set and showed similar reconstruction loss (Figure S4G).

The VAE latent space also captured several other key features of PRC2 control of the developing CNS. These included the graded involvement of PRC2 along the A-P axis, with extensive gene up-regulation in the *Eed-cKO* FB and smaller effects in the HB (Figure 4D). We also observed a temporal reduction in the H3K27me3 mark, as evident in the MB from E10.5 to E16.5 (4D).

The posterior gene cohort was repressed in the FB and MB in WT and upregulated in *Eed-cKO* (Figure 5A). This cohort was enriched for ChromHMM bivalent promoter states, suggesting that these genes are directly controlled by PRC2 and are selectively expressed (Figure 5A). The anterior gene cohort tended to exhibit an opposing RNA expression profile to the posterior genes, with a decrease in expression over time, and limited enrichment of H3K27me3-associated chromatin states (Figure 5B). The development cohort included a mixture of genes that were mostly upregulated in *Eed-cKO*, and whose ChromHMM profile indicated both direct and indirect PRC2 effects (Figure 5C). The unmarked proliferation cohort was enriched for cell cycle genes, mostly those with pro-proliferative function (Figure 5D). In WT, genes in this cohort displayed a logical downregulation as neurogenesis comes to an end in both FB and SC. Relative to WT, PRC2 inactivation accelerated the decrease in expression of this cohort in all tissues, but most distinctly in FB (Figure 5D). Lastly, the immune response cohort was weakly upregulated in *Eed-cKO* and enriched for the absence of PRC2-associated ChromHMM marks, indicating an indirect effect of PRC2 (Figure 5E).

**Fig. 5.**
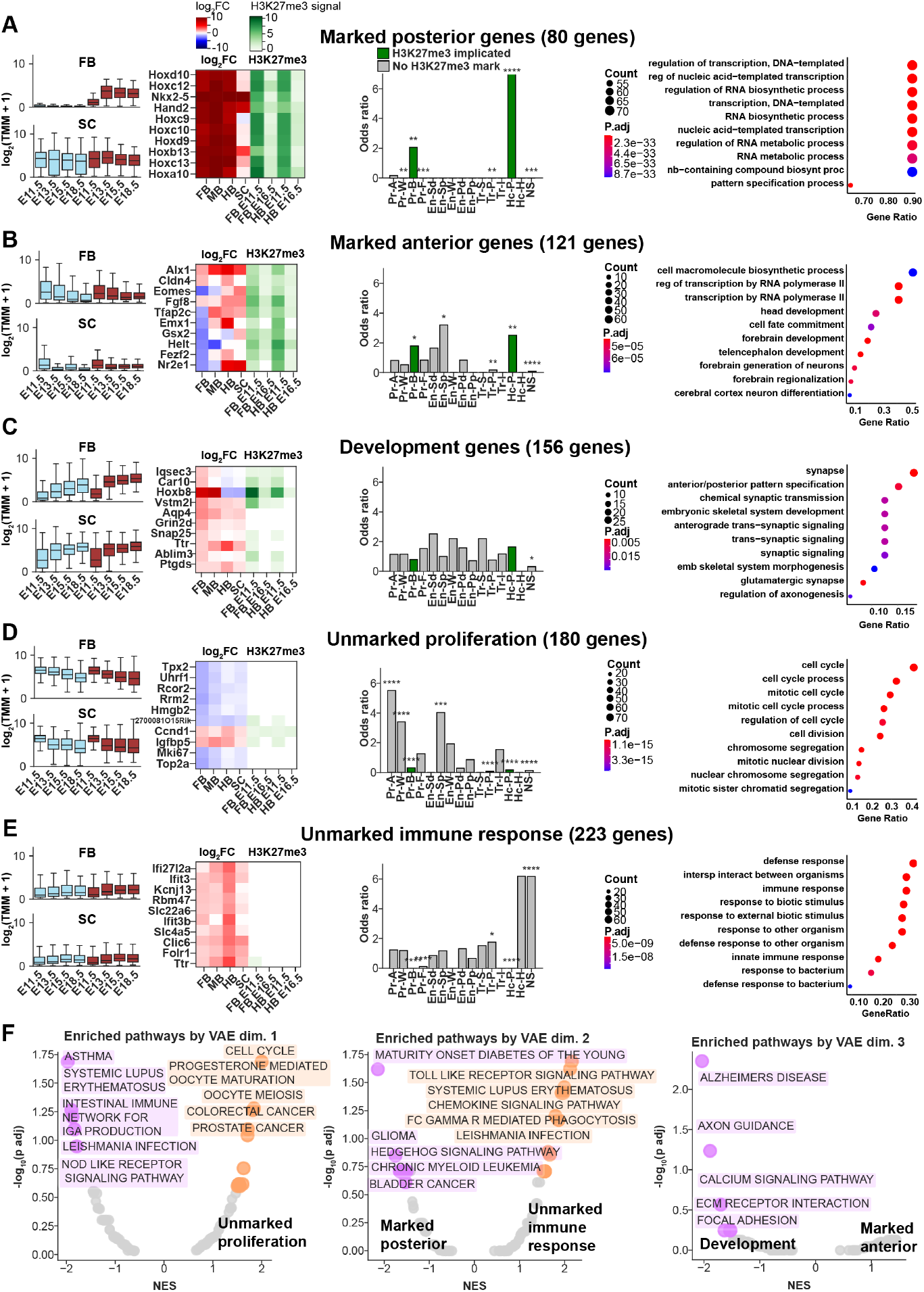
VAE identifies functionally diverse gene cohorts. (A-E) Different “cohorts” of genes identified at tails of one or more VAE latent dimensions (+ 1.25 SD). (A) In the WT, the marked posterior gene cohort is repressed in the FB and expressed in the SC. In the *Eed-cKO* mutants, they are overexpressed in FB, while the SC is unaffected. Top-10 genes in this group are predominantly Hox genes, marked by H3K27me3 in both FB and HB. Enrichment of ChromHMM chromatin states and GO terms show bivalent and repressive states, and gene regulation, respectively. (B) In WT, the marked anterior gene cohort is repressed in SC and expressed in FB. In the *Eed-cKO* mutants, they are mostly downregulated in FB and upregulated in SC. Top-10 genes display tissue specific response to *Eed-cKO*. They show no enrichment of specific chromatin states, but FB differentiation and development GO terms. (C) The development gene cohort increases in expression over time and is more highly expressed in SC than FB, a trend that is more pronounced in the mutant. Top-10 genes display upregulation in the mutant and are both marked and unmarked. Enrichment analyses show no enrichment of specific chromatin states, but GO terms for embryonic development. (D) The unmarked proliferation gene cohort decreases in expression over time, in both the WT and *Eed-cKO*, which is pronounced in *Eed-cKO*. These genes are enriched for active and weak promoter chromatin states and cell cycle functions, suggesting that active genes are important for cell growth and the rate of proliferation. (E) The unmarked immune response gene cohort displays no specific expression profile in WT, but are upregulated in the mutant. Top-10 genes show a homogenous response to *Eed-cKO*, in particular a strong upregulation in HB. These genes are enriched for the no signal (NS) ChromHMM chromatin state. Enrichment for defence and immune response GO terms indicates functional homogeneity. (F) Genes ordered by each VAE dimension enrich in KEGG pathways (top-5 shown, plotted by normalised enrichment score (NES) v. transformed P-value), which concord with GO analysis and reinforce the characterisation of each gene cohort, e.g., unmarked proliferation genes map to large tail of dimension 1, and intersect with pathways for “cell cycle” and “oocyte meiosis”.

### VAE latent dimensions identify allied but functionally diverse genes

The variation captured by the VAE enabled the discovery of new categories of genes that were coinciding in each latent dimension. We grouped genes by their existence at the extremes of each dimension, with membership determined by being in the tail of the distribution (*SD* > 1.25 from mean), resulting in six non-exclusive cohorts of genes. One cohort was omitted from further analysis, as it contained genes that extensively overlapped with the other five cohorts (Figure S11). For the five remaining cohorts we used GO term enrichment and selective gene expression to manually label the cohorts, yielding: (1) posterior genes, (2) anterior genes, (3) development genes, (4) unmarked proliferation genes, and (5) immune response genes (Figure 5A-E).

The posterior gene cohort was repressed in the FB and MB in WT and upregulated in *Eed-cKO* (Figure 5A). This cohort was enriched for ChromHMM bivalent promoter states, suggesting that these genes are directly controlled by PRC2 and are selectively expressed (Figure 5A). The anterior gene cohort tended to exhibit an opposing RNA expression profile to the posterior genes, with a decrease in expression over time, and limited enrichment of H3K27me3-associated chromatin states (Figure 5B). The development cohort included a mixture of genes that were mostly upregulated in *Eed-cKO*, and whose ChromHMM profile indicated both direct and indirect PRC2 effects (Figure 5C). The unmarked proliferation cohort was enriched for cell cycle genes, mostly those with pro-proliferative function (Figure 5D). In WT, genes in this cohort displayed a logical downregulation as neurogenesis comes to an end in both FB and SC. Relative to WT, PRC2 inactivation accelerated the decrease in expression of this cohort in all tissues, but most distinctly in FB (Figure 5D). Lastly, the immune response cohort was weakly upregulated in *Eed-cKO* and enriched for the absence of PRC2-associated ChromHMM marks, indicating an indirect effect of PRC2 (Figure 5E).

To address if the WT profile of gene cohorts were conserved in humans, in particular regarding the expression of the genes in the marked anterior cohort, we analysed publicly available data from PsychEncode (56). We found similar expression patterns across the mouse and human orthologs, with a significantly greater expression in human FB tissue of the genes we identify to be FB specific, than in other brain tissues (Figure S10).

### VAE uniquely recovers allied genes in A-P axis development

Similar to its “deep” neural network cousin, the VAE allows low-dimensional codes to capture non-linear relationships from a high-dimensional input space, staggered at each intermediate layer (57). To quantify the level of organisation of the VAE relative to other dimensionality methods: PCA, UMAP, tSNE, and PHATE, we performed a number of tests.

We asked if each method at *D* = 3 had the capacity to find latent codes that distinguished marker genes by their known A-P association. Only the distances in the VAE projection were able significantly distinguish between marker gene sets in both the consistently affected and partly affected datasets (Figure S5A-B).

The insights reported in previous sections were based on five gene cohorts that were evident in VAE at *D* = 3. We next confirmed that VAE reproduced similar biological meaning at *D* = 6, and asked if alternative methods were able to extract groups also with similar biological meaning. Finally, each method was used to select the tailing 200 genes in each latent component at D=3, these gene groups were then tested for functional enrichment. All methods were able to extract the most salient functional groups, e.g., the posterior group. However, only the VAE and tSNE were able to distinguish an anterior gene cohort (Figure 5B, S8). PCA and UMAP contained duplications in encoding for the cell cycle (Figure S6, S7), while PHATE uniquely identified a group containing “membrane” and “signalling” terms (Figure S9). While tSNE functions at a comparable level to the VAE in terms of pathway and GO enrichment, it was unable to separate gene sets when used as a distance metric (Figure S5).

### PRC2 regulates cell cycle genes directly or by proxy TFs

A key phenotype of *Eed-cKO* mutants is a striking reduction of proliferation of the FB (Figure 1I-J) ((5)). Moreover, the VAE analysis identified many genes (180 genes), in the “unmarked proliferation” cohort (5D). These findings prompted us to focus on the expression of the main cell cycle genes (32 genes; Table S1), to explore the regulation of pro- and anti-proliferative genes. Looking first at WT, we observed that the majority of pro-proliferative and anti-proliferative genes were expressed in opposing gradients along the A-P axis (Figure 6A, D). In *Eed-cKO* mutants, we found that the majority of cell cycle genes (29/32) were affected, with 8 consistently affected and 21 partly affected (Figure 6). With few exceptions, pro-proliferative genes were downregulated while anti-proliferative genes were upregulated relative to WT, and these effects were most pronounced in the anterior CNS (FB and MB), resulting in a general flattening of the gene expression gradients (Figure 6A, 6D). Two outliers were the pro-proliferative genes *Ccnd1/2*, which were strongly upregulated in the posterior CNS (HB and SC) (Figure 6A). The H3K27me3 profiles of the anti-proliferative genes were comparatively pronounced, although several pro-proliferative genes, such as *Ccnd1/2* and *Ccna1*, were also marked (Figure 6A).

**Fig. 6.**
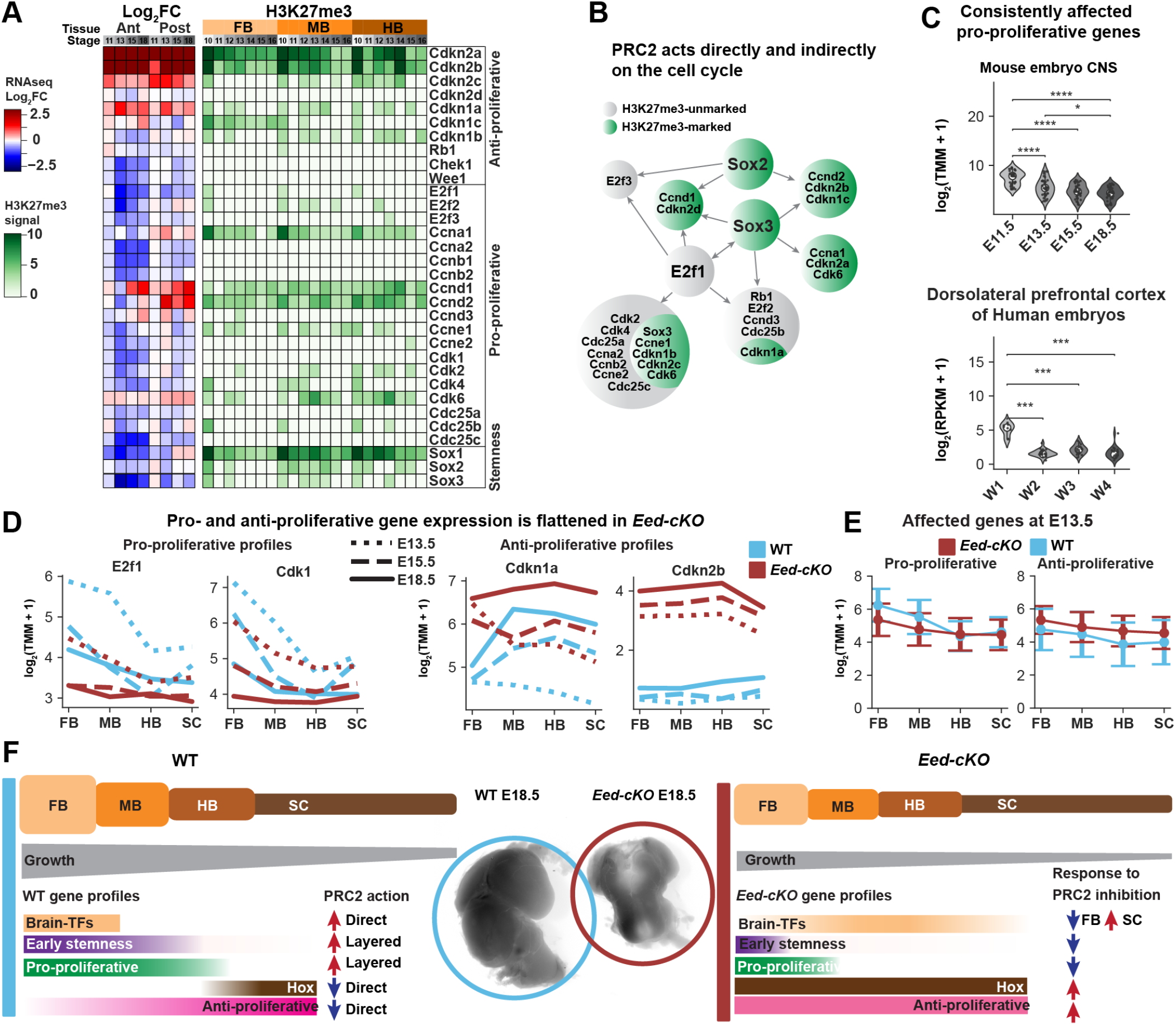
Layered cell control by PRC2. (A) Cell cycle gene response to PRC2 inactivation shows strong upregulation of some marked and consistently affected anti-proliferative genes, while the majority of pro-proliferative genes are downregulated and unmarked. Sox genes are marked and downregulated. (B) Proposed mechanism of action for the indirect regulation of the Sox genes on the cell cycle genes. (C) Consistently affected pro-proliferative genes (E2f1, Ccna2, Ccnb1, Cdc25c and Ccnd1) exhibit a reduction in expression over embryonic development. This trend is observed in mice across the brain (FB and MB), and in human embryonic samples, also across the brain (dorsolateral prefrontal cortex). (D) Select cell cycle genes exhibit evidence of an A-P gradient in WT. (E) Grouping all affected cell cycle genes reveals a trend (mean and standard error) for to A-P flattening in expression. (F) PRC2 ensures that Hox homeotic genes are only expressed in the SC and HB, and brain TFs only in the FB and MB, and promotes gradients of stemness, anti- and pro-proliferative gene expression. These A-P differences in gene expression drives an earlier progenitor proliferation stop and more limited daughter cell proliferation in the posterior CNS, creating a gradient of growth.

Because the majority of pro-proliferative genes appeared to be indirectly affected by PRC2, we sought to identify which TFs could be targeting the proliferation genes. We again focused on the *Sox1/2/3* stemness genes, as well as E2f1, a core TF in the cell cycle machinery (58). *Sox1/2/3* were marked by H3K27me3, while *E2f1* showed little if any marks (Figure 6A). However, all four genes were downregulated in *Eed-cKO* (Figure 6A). Previous ChIP-seq studies have probed the genome-wide occupancy of three of these four TFs (59, 60, 61). These data revealed that Sox2 and −3 bind to a number of proliferation genes, including E2f1 and other E2f genes, and that E2f1 binding showed extensive overlap with the Sox2/3 binding profiles (Figure 6B). These findings suggest that PRC2 action is layered – acting both directly and indirectly, via *Sox1/2/3* and *E2f1/2/3*, to control cell cycle gene expression.

To investigate whether the cell cycle gene expression profiles are evolutionarily conserved, we tested whether the WT profile of early activation of pro-proliferative genes is conserved in humans. We again used the publicly available data from PsychEncode (56) and confirmed a significant reduction over time in the pro-proliferative genes in human embryonic brain development (Figure 6C).

## Discussion

### PRC2 promotes the developing CNS A-P axis

The developing CNS displays a striking and evolutionarily conserved patterning along the A-P axis, evident by the selective expression of brain-specific TFs anteriorly and Hox homeotic genes posteriorly (62, 63). Studies in *Drosophila* have also revealed an A-P expression gradient of neural stemness genes e.g., the SoxB family (2). In *Drosophila*, the selective expression of brain-TFs, Hox genes and neural stemness genes is accompanied by and (to a great extent) drives gradients in pro- and antiproliferative gene expression, which in turn results in a gradient of progenitor and daughter cell proliferation, faster cell cycles, and the expansion of the anterior CNS (2, 3, 4, 5). Studies in mouse have indicated that many of these developmental features are conserved in mammals, although the degree of conservation is unclear (5). Moreover, while PRC2 plays a key role in promoting these A-P differences, its precise roles have hitherto not been comprehensively addressed.

We created a transcriptomic dataset of the WT and *Eed-cKO* mutant developing mouse CNS, covering the major phase of neurogenesis. We observed the anticipated WT expression of brain-TFs and Hox genes, anteriorly and posteriorly, respectively. In addition, we observed striking gene expression gradients of stemness, pro- and anti-proliferative genes, demonstrating that these features are also conserved from *Drosophila* to mouse. We found that PRC2 inactivation resulted in profound gene expression changes in the CNS, which are particularly pronounced in the anterior CNS, with the FB displaying 4,414 DE genes compared to 771 DE genes in the SC. Looking specifically at the aforementioned developmental genes we found that PRC2 inactivation reduced brain-TF expression and upregulated Hox genes anteriorly. In addition, we observed a flattening of the gene expression gradient of stemness and pro-proliferative genes, and an upregulation of anti-proliferative genes. Hence, PRC2 plays a fundamental role in promoting anterior CNS development, with anterior tissues posteriorizing and reducing their stemness in *Eed-cKO* mutants (Figure 6F). These regulatory effects generally accumulate over time i.e., once a gene becomes dysregulated it remains so, and hence the maximum difference in DEGs between WT and *Eed-cKO* is at E18.5, the latest stage sampled herein.

### PRC2 inactivation causes extensive direct and indirect effects

To understand if PRC2 acts in a direct or indirect manner upon the affected genes, we integrated our transcriptomic dataset with histone modification profiles, generated by ENCODE. This label-stratified analysis identified six gene categories, based upon genes being H3K27me3-marked or not, and upon genes having expression levels that are unaffected, partly or consistently affected.

All partly/consistently affected genes with H3K27me3 (2,494 combined) were enriched for GO terms related to embryonic patterning, which aligns well with the observed effect of *Eed* mutation i.e., a flattening of the CNS A-P axis. This finding, combined with their ChromHMM states, indicates that this gene group is directly regulated by PRC2. The category of H3K27me3-marked and unaffected genes (410 genes) was enriched for similar GO terms i.e., regulation and development, showing that a subset of H3K27me3-marked developmental genes are not affected by *Eed-cKO*.

There were many partly/consistently affected genes without H3K27me3 (10,525 genes combined), suggesting an extensive indirect effect of PRC2 inactivation. The consistently affected genes primarily included immune response genes (many of which were also identified by the VAE analysis, see below), while the partly affected category included RNA processing genes.

### VAEs distinguish allied gene cohorts relevant to A-P axis control

To further address the function of PRC2 and to tease apart the layered roles of PRC2 we used a deep learning approach. To this end, we applied a VAE to exhaustively probe gene expression and H3K27me3 patterns along A-P development, over several timepoints, in control and PRC2 mutant. The VAE was able to distinguish between cohorts of genes with qualitatively different functional profiles and multi-variate trends across the datasets. Moreover, despite sharing at least one latent label, several genes within each cohort were surprisingly varied in terms of both expression changes and chromatin state, indicating that the multi-variate nature of the VAE analysis uncovers a spectrum of biologically relevant, gene groupings across gene expression and histone modification features. While we could have extended our label stratified analysis to include other factors, e.g., “up” or “down” in each DE analysis, the number of gene categories increases exponentially. In contrast, the VAE identified functionally enriched gene groups important for CNS development using only three dimensions.

When comparing the VAE analysis to the initial, label-stratified analysis, the VAE identified additional gene cohorts and was able to tease apart the functional roles identified within the label-stratified categories. This includes the separation of the “marked, consistently affected” group, containing both anteriorly and posteriorly expressed genes, into the “marked anterior” and “marked posterior” cohorts. When comparing the VAE to other dimensionality reduction methods we found that only the VAE and tSNE recovered functionally enriched anterior and posterior groups. However, the VAE methods were most accurate at distinguishing between “like” and “unlike” gene sets both the consistently and partly affected datasets. We developed our analysis workflow and package to be applicable to other biological domains. In particular, this analysis is amenable to any system where a biological features profile (e.g. representing the state of a gene) across an assay of experiments is indicative of function and mode of regulation. For example, to identify distinct regulatory responses between patients cohorts in tumours, understanding TF activation through development, or regulatory responses to drugs.

### Immune response genes may be affected by several mechanisms

Unexpectedly, both the label-stratified and VAE analyses identified immune response genes as a salient function affected by PRC2. Using *Sox1-Cre* to delete *Eed* only removes gene function in the CNS itself, and not in the blood or blood vessels (Figure 1E; (5)). It is therefore possible that the undergrown FB and MB in *Eed-cKO* mutants resulted in a higher ratio of blood vessels/immune cells to CNS cells, thereby increasing the transcriptome signal for immune response genes in an indirect manner. However, two other plausible causes of activation of immune response genes are (1) a CNS-autonomous effect, as PRC2 has been linked to immune responses in human cancer (64) and/or (2) that the developmental defects in *Eed-cKO* mutants lead to a breakdown of the blood brain barrier and/or an immune response to a malforming CNS. Further studies i.e., spatio-temporal single cell RNA-seq, would be required to determine why the immune response genes are activated.

### Multi-layered control of proliferation by PRC2

While the label-stratified analysis identified many cell cycle genes, they were distributed across several gene categories. In contrast, the VAE grouped them into a consistent gene cohort: the proliferation cohort. In general, pro-proliferative genes were downregulated and anti-proliferative genes upregulated, and there was a general flattening of their A-P expression gradients. These gene expression changes are likely directly responsible for the undergrowth phenotype observed in the mutant FB and MB. Analysis of the H3K27me3 profiles revealed that PRC2 may be acting directly on a subset of marked proliferation genes, and likely indirectly, via e.g., the Sox1/2/3 and E2f1/2/3 TFs, on un-marked proliferation genes.

The tendency for PRC2 to directly regulate anti-proliferative genes and indirectly regulate pro-proliferative genes, points to an uneven involvement of the epigenetic machinery in cell cycle regulation. This finding is not surprising given the different evolutionary age of the cell cycle genes and the gradual emergence of the epigenetic machinery. Specifically, while the basic core cassette of cyclins and Cdks is ancient in eukaryotes (65) the Kip/Cip family evolved later, and INK4 even more recently (the INK4 family is not present in *Drosophila*). The Kip/Cip and INK4 families likely evolved to provide the increasingly refined control of proliferation necessary in larger metazoans. Indeed, evolution of the cell cycle machinery has gone hand in hand with, and one may argue been facilitated by, an increasingly elaborate epigenetic machinery. Against this backdrop, it is logical that PRC2 is heavily engaged in directly regulating the Cip/Kip and INK4 families, but indirectly regulating the ancient cell cycle genes.

### PRC2 gates an ancient CNS stemness gradient

One of the key features of the developing CNS A-P axis is a stemness gradient, which drives CNS anterior expansion. PRC2 plays five key roles herein: (1) promoting brain-specific TF expression, (2) repressing anterior Hox gene expression, (3) promoting a gradient of neural stemness TF expression, (4), repressing anterior anti-proliferative gene expression and, (5) promoting anterior pro-proliferative genes (1). Our findings herein suggest that PRC2 regulates the first four categories directly by application of H3K27me3; PRC2 regulates pro-proliferative genes by also relying on proxy TFs.

Our spatio-temporal transcriptomic and epigenomic analysis provides an in-depth view into the strikingly different regulatory landscape present in the anterior versus posterior regions of the CNS, and the profound importance of PRC2 in establishing and driving these differences. Previous studies show that the role of PRC2 in gating A-P gene expression is integral for mouse, fly, and zebrafish development (8, 66, 67). Our work extends upon this, revealing that the FB genes dysregulated in the developing mouse PRC2 mutant CNS are also selectively expressed during human FB embryonic development, underscoring the evolutionary conservation of brain development across bilateria.

A number of observations in different species, including gene expression analysis, and anatomical and phylogenetic considerations, have led to the proposal that the anterior and posterior CNS may have originated from different parts of the nervous system present in the Bilaterian ancestor, the apical and basal nervous systems (62, 68, 69, 70). If true, this brain-nerve cord “fusion” concept may help explain the strikingly different gene expression and neurogenesis properties of the brain, when compared to the nerve cord, as well as the apparent “brain-preoccupation” of the PRC2 complex.

### Data and code availability

Raw RNA-seq files are available at the NCBI/Gene Expression Omnibus under the accession GSE123331. Python code including Jupyter Notebooks (both as HTML and ipynb) used to generate all results are available at: https://arianemora.github.io/mouseCNS_vae/

## Lead contacts

Further information and requests for experimental protocols should be directed Stefan Thor, information and requests for computational methods should be directed to Mikael Bodén.

## ACKNOWLEDGEMENTS

We are grateful to Jose Dias, Johan Ericson, The Jackson Laboratory mouse stock centre and the ENCODE consortium, for sharing reagents and advice. We thank Tyrone Chen and Don Teng for critically reading the manuscript. Carolin Jonsson and Helen Ekman provided excellent technical assistance.

## AUTHOR CONTRIBUTIONS

A.M, M.B and S.T designed the analysis and A.M implemented the software. J.R, S.T designed the experiments and S.T, J.R, I.M.C, B.Y and A.S conducted the experiments. A.M, M.B and S.T wrote the manuscript. All authors reviewed the submitted material.

## Supplemental information

**Supplemental Table 1.**
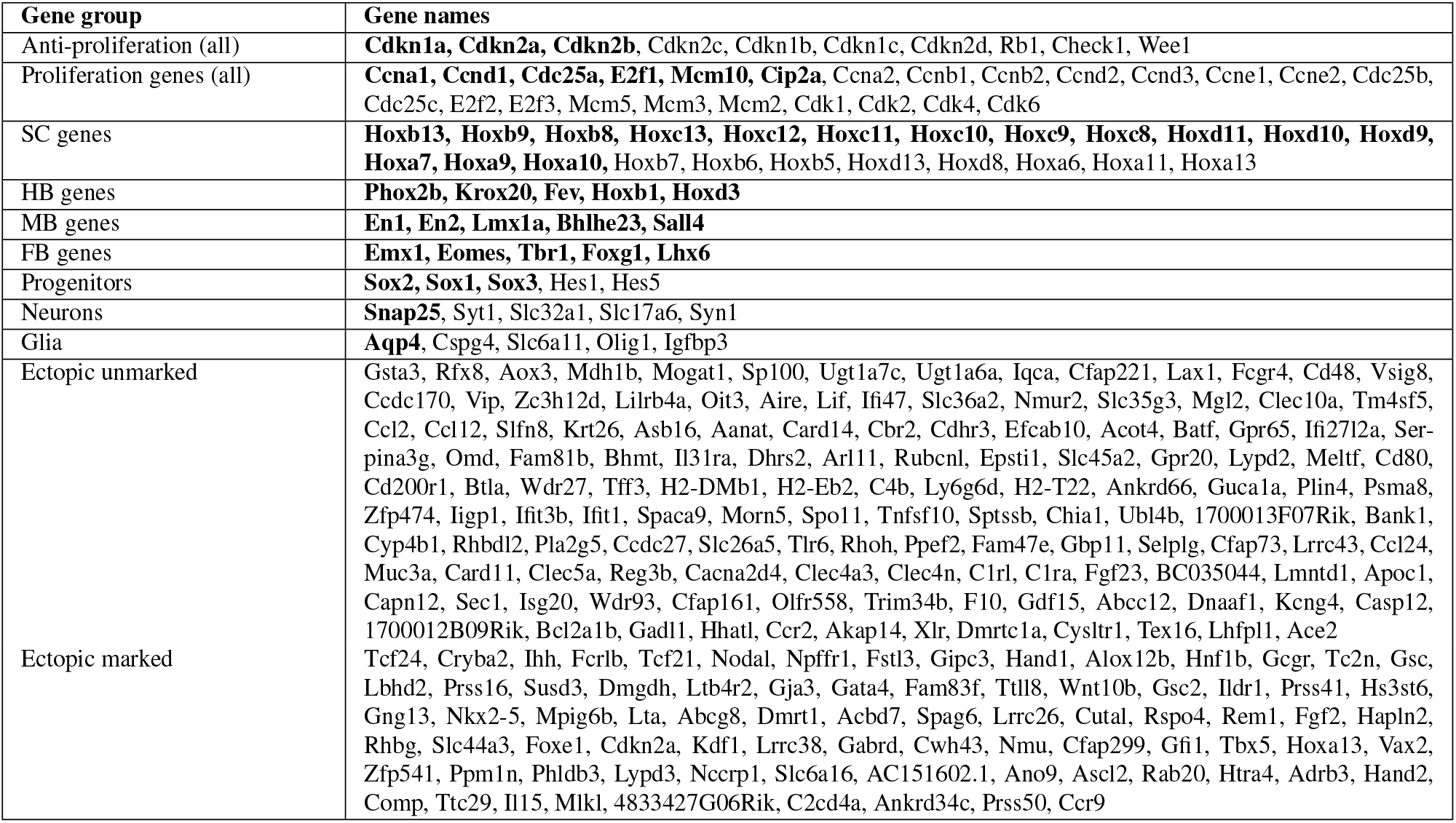
Gene lists Genes sets used throughout the paper, bolded genes were used for display in Figure 4. Display genes were chosen based upon published gene expression patterns. Ectopic genes were identified computationally and indicates that the gene had a mean expression less than 0.5 TMM in the WT and greater than 0.5 TMM in the mutant.

**Supplemental Table 2.** Normalised SSQ distance between tissues For each *Eed-cKO* condition the distance (normalised sum of square differences between gene expression) between the mutant’s merged replicates and all WT conditions is shown. The WT condition with the smallest distance is used. There is an evident regression along the A-P axis of FB samples, and some MB samples (highlighted in bold).

| *Eed-cko* | Most similar WT condition | Distance |
| --- | --- | --- |
| ko11fb | wt11mb | 32.689 |
| ko13fb | wt13mb | 67.778 |
| ko15fb | wt15mb | 60.334 |
| ko18fb | wt18hb | 66.041 |
| ko11mb | wt11hb | 29.972 |
| ko13mb | wt13sc | 56.335 |
| ko15mb | wt15mb | 54.135 |
| ko18mb | wt18hb | 57.589 |
| ko11hb | wt11hb | 36.108 |
| ko13hb | wt13sc | 58.831 |
| ko15hb | wt18hb | 60.739 |
| ko18hb | wt18hb | 54.093 |
| ko11sc | wt11sc | 24.458 |
| ko13sc | wt13sc | 37.104 |
| ko15sc | wt18sc | 51.529 |
| ko18sc | wt18sc | 39.241 |

**Supplemental Fig 1.**
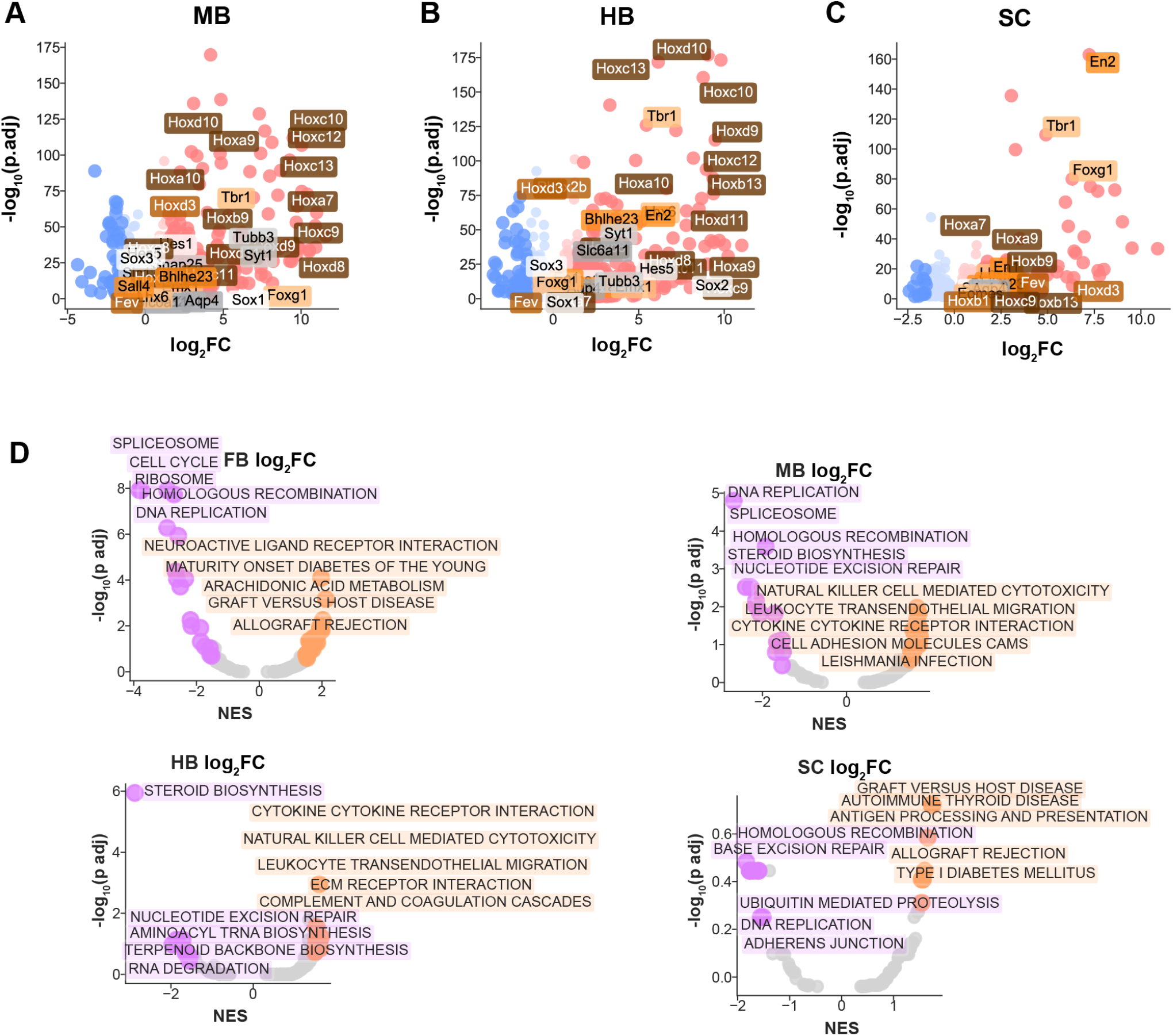
Tissue specific response to *Eed-cKO*. (A) In MB Hox genes are upregulated in Eed-cKO for E13.5 – E18.5 as show by a standard Volcano plot (log-fold change v. log P-value). (B) In HB Hox genes and select FB genes, such as Tbr1, are upregulated in Eed-cKO, plotted by normalised enrichment score (NES) v. transformed P-value. (C) In SC FB specific genes upregulated in Eed-cKO, with an overall smaller response than in MB and HB. (D) Using the logFC for each tissue to rank the significant genes for the given experiment, in FB and MB there is negative enrichment for the cell cycle pathway and RNA biology pathways, such as “ribosome” and “spliceosome” (these do not appear to be negatively enriched in the HB and SC). In all four tissues the top positively enriched pathways are associated with immune response. In the posterior tissues there is negative enrichment of some overlapping pathways with the anterior tissues, such as “DNA replication”.

**Supplemental Fig 2.**
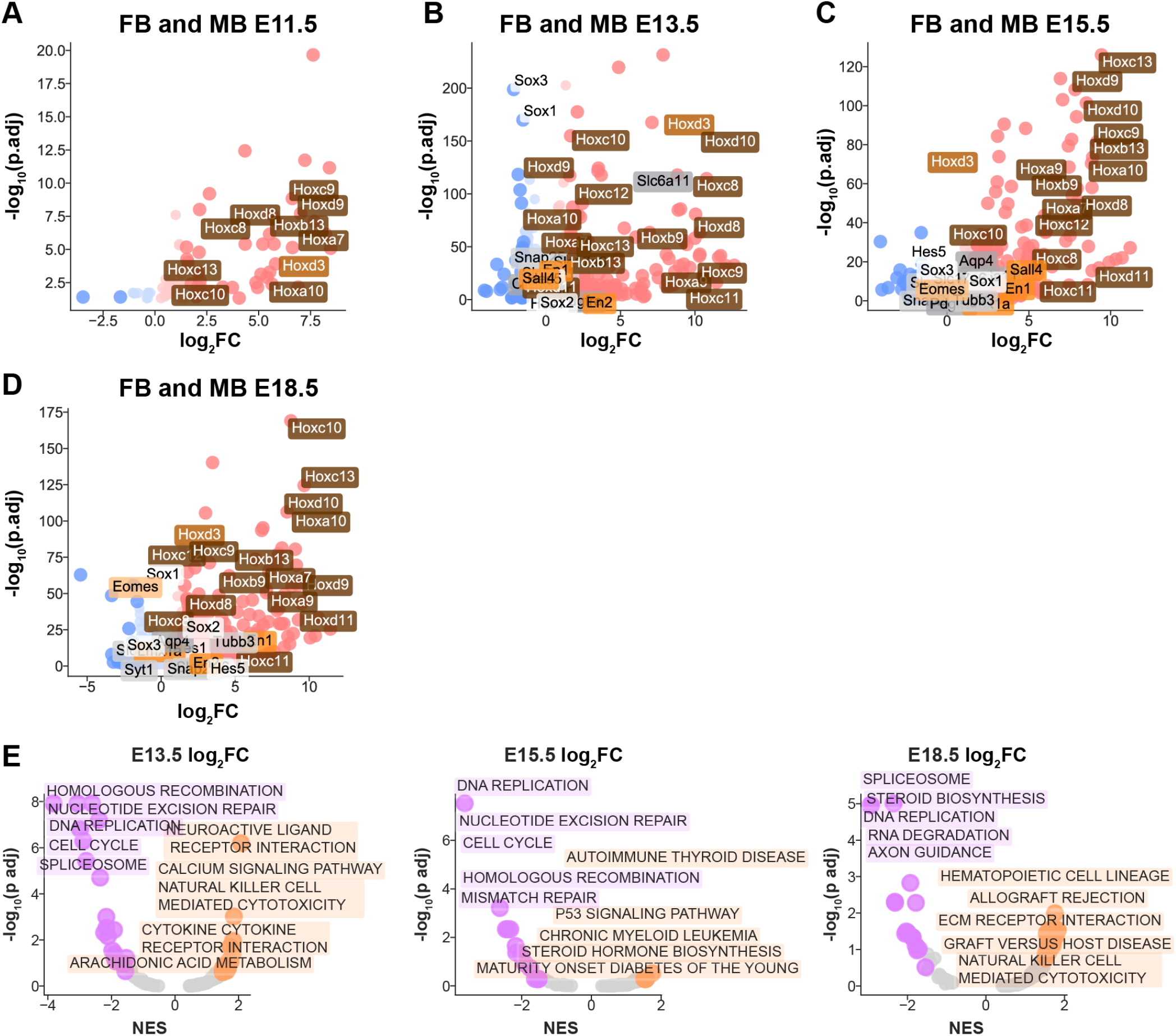
Temporal response to *Eed-cKO* in anterior CNS. (A) Anterior (FB, MB) regions at E11.5 shows limited upregulation of Hox genes and overall minor effects. (B) At E13.5, there is strong downregulation of progenitor markers *Sox3* and *Sox1* and upregulation of Hox genes. (C) At E15.5 there are fewer downregulated genes, however displaying a stronger upregulation response, in particular of Hox genes. (D) At E18.5 there is a similar upregulation as the earlier time points of Hox genes and strong downregulation of forebrain marker, *Eomes*. (E) At E13.5 and E15.5 there is downregulation of the “cell cycle” pathway. At E18.5 “cell cycle” is no longer a significant term but other terms such as “DNA replication” are shared. In the positively enriched pathways there is enrichment of immune associated pathways, and a response to a disrupted system (e.g., “allograft rejection”)

**Supplemental Fig 3.**
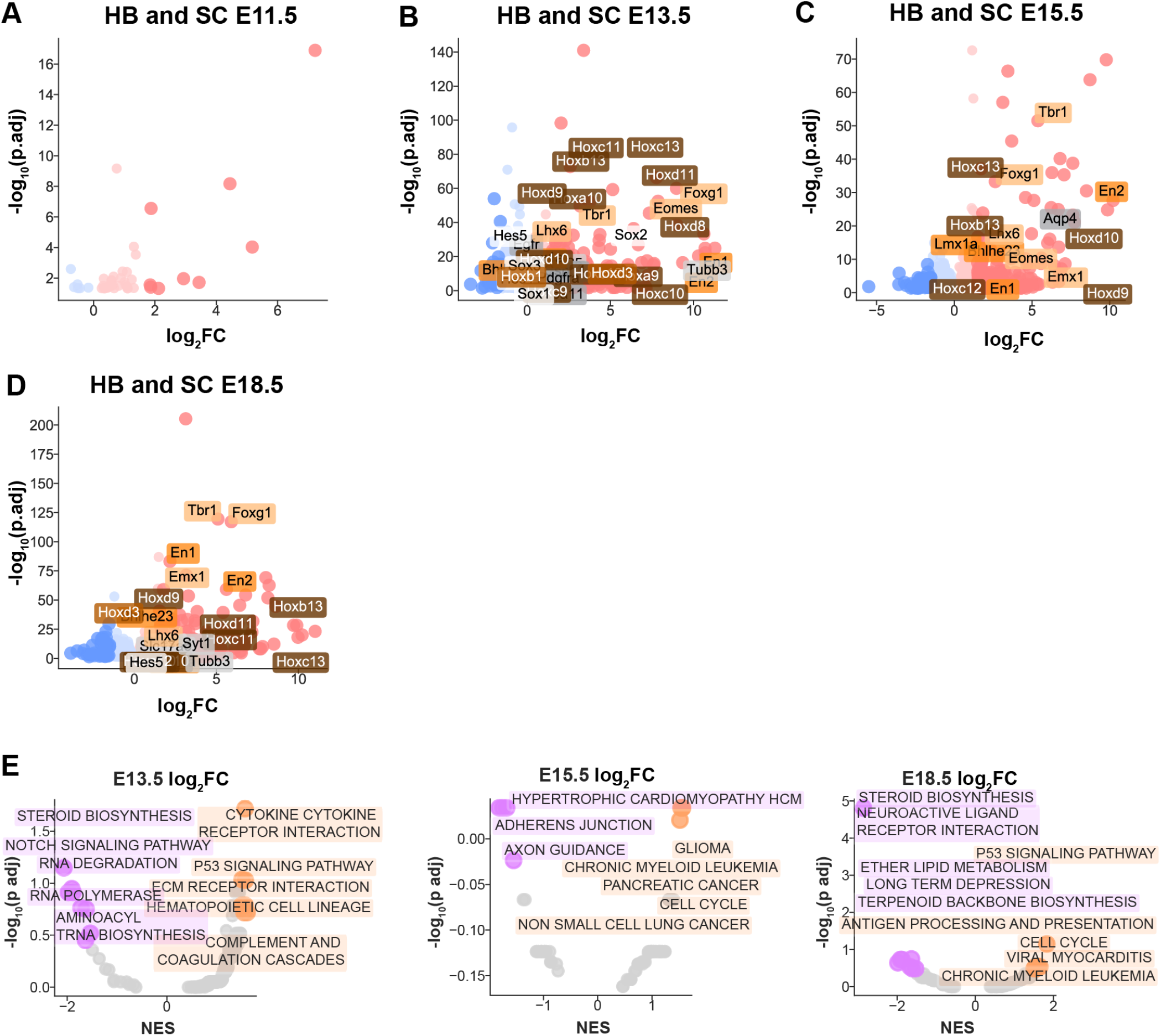
Temporal response to *Eed-cKO* in posterior CNS. (A) The logFC response in the posterior (HB, SC) regions at E11.5 shows minimal effects in Eed-cKO. (B) At E13.5 there is upregulation of both SC and FB markers. (C) At E15.5 there is upregulation of markers from across brain regions. (D) At E18.5 the same effects of ubiquitous upregulation are observed. (E) At E13.5 there is a downregulation of RNA pathways (“RNA degradation”, “RNA polymerase”), and an upregulation of immune response or cancer terms, as well as of “blood cells” and “hematopoietic cell lineage”. At E15.5 and E18.5 there is upregulation of the cell cycle pathway and also of blood cancer associated pathways, such as “chromic myeloid leukemia”. At the later stages there is negative enrichment for brain associated pathways, such as “neuroactive ligand receptor interaction” and “axon guidance”.

**Supplemental Fig 4.**
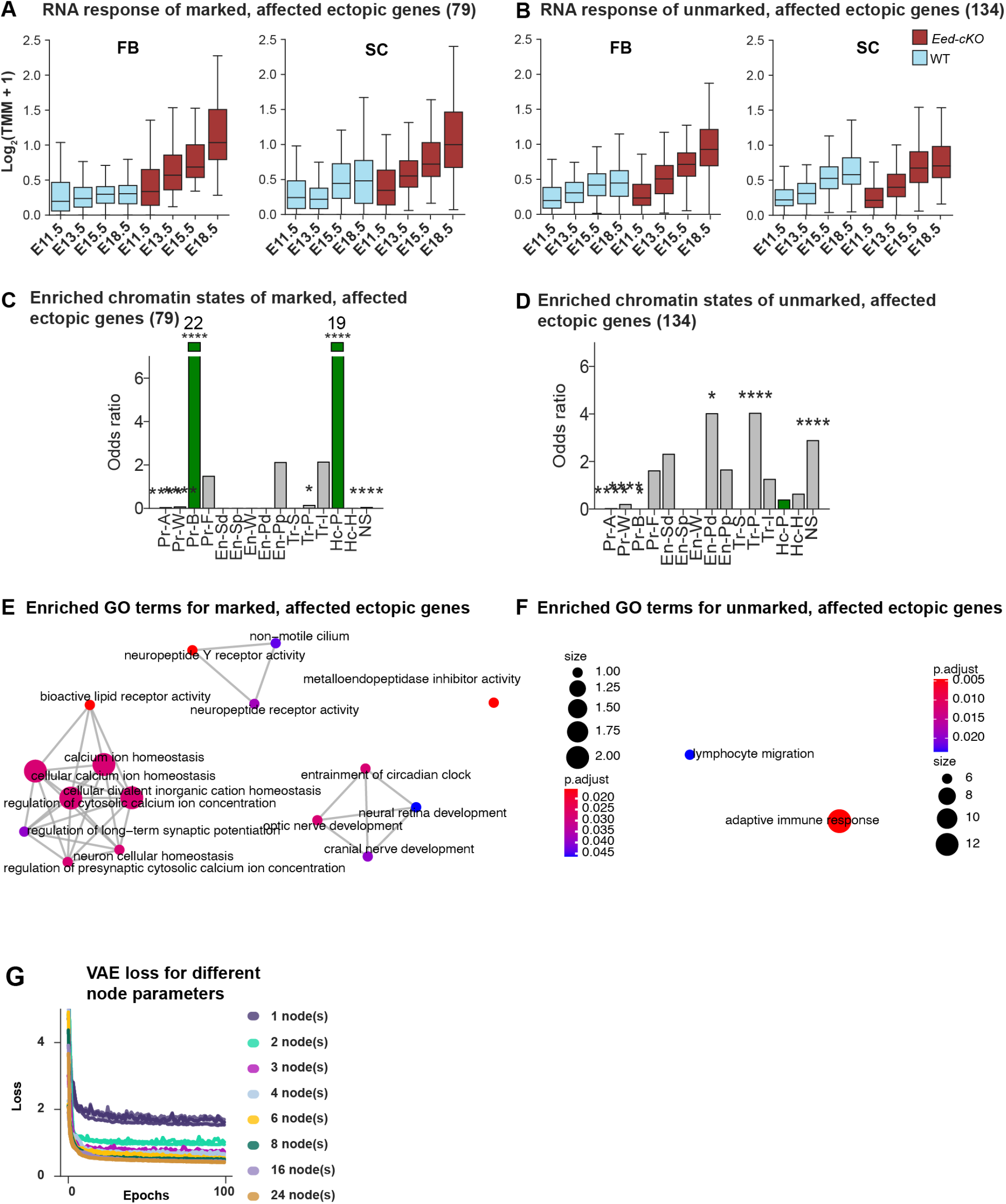
Ectopically expressed peripheral genes in *Eed-cKO*. (A) Marked peripherally expressed genes that were ectopically expressed in Eed-cKO, are upregulated over time in both FB and SC. (B) Unmarked peripherally expressed genes that were ectopically expressed in Eed-cKO are upregulated over time in FB, but with limited effects in SC. (C) Marked genes are enriched for ChromHMM signature of bivalency, while (D) unmarked genes are enriched for no signal (NS), and permissive transcription state (Tr-P). (E) GO analysis of marked genes shows a diverse range of terms. (F) Few terms were associated with the unmarked group. (G) Loss of the VAE on the 1,371 dataset shows the loss stabilises at 3 latent dimensions, with marginal improvements for greater numbers.

**Supplemental Fig 5.**
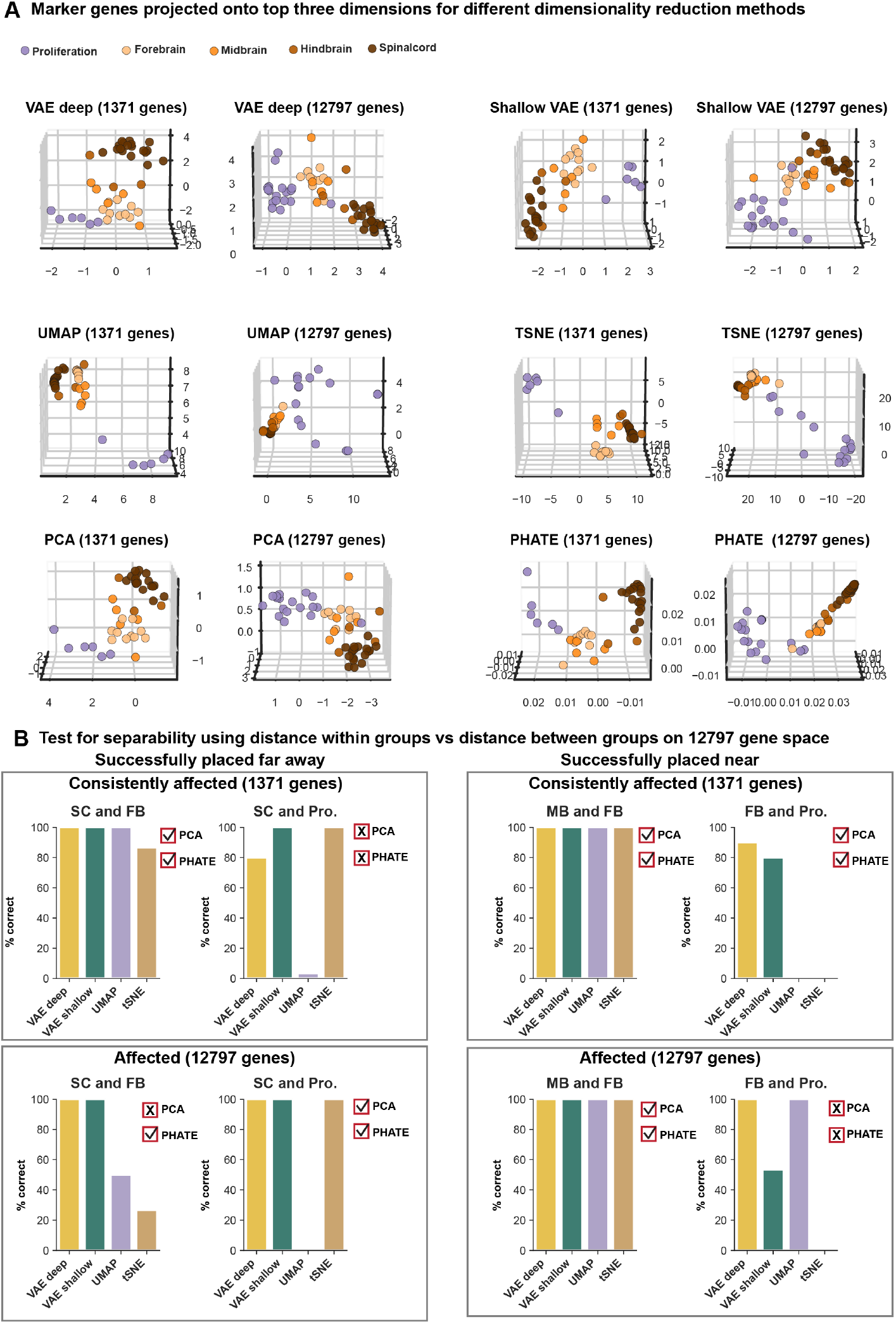
Marker genes projected into alternate spaces. (A) Marker genes were projected onto 3-dimensional latent spaces produced by five methods: PCA, tSNE, UMAP, PHATE and a shallow VAE (a VAE with only one internal layer). Separability between FB, MB, HB, and SC is most apparent when using the consistently affected gene set (1,371 genes). Separability is reduced with most tools when using the larger dataset (all affected genes). The VAE and PCA appear most robust to adding partly affected genes. (B) Testing separability between the most diverse gene groups over 30 runs for non-deterministic tools (tSNE, VAE, UMAP), and from deterministic tools PCA and PHATE revealed that VAE methods were most accurate at assigning small distances within a marker group and larger distances between marker groups (i.e. large distance within a cluster when combining un-related gene groups).

**Supplemental Fig 6.**
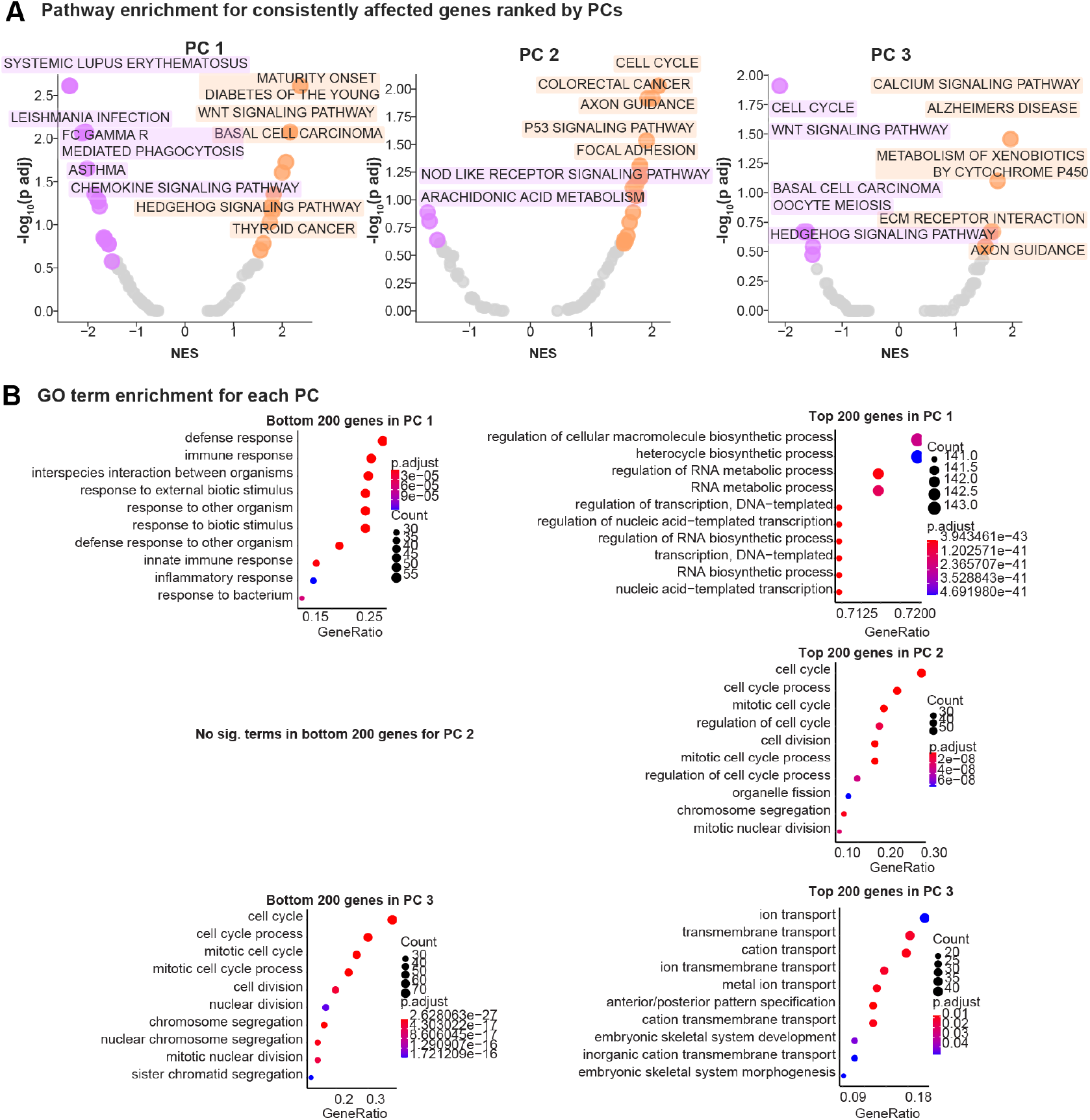
Enrichment for PCA. (A) Genes were ranked by each Principal Component (PC) (up to 3) to identify negatively and positively enriched pathways. PC 1 is negatively enriched for immune associated pathways, and positively enriched for brain dysregulation, “WNT signalling, Hedgehog signalling” and cancer pathways. PC 2 is negatively enriched for only two pathways: “Nod like receptor” and “arachidonic acid metabolism”, which is associated with neurotransmitter systems. The top five positive pathways in PC 2 are diverse, including “cell cycle”, “axon guidance” and cancer pathways. PC 3 is negatively enriched for diverse pathways, including cell cycle, carcinoma, and Hedgehog signalling. PC 3 is positively enriched for brain associated pathways, including “Alzheimers” and “axon guidance”. (B) The top and bottom 200 genes along each PC were tested for enriched GO terms. PC 1 agrees with the enrichment of pathways, e.g., with RNA metabolism associated GO terms positively enriched (contain Hox genes). In PC 2 the top 200 genes are not enriched for any GO terms, while the bottom 200 enriched predominately for cell cycle terms. PC 3 is negatively enriched for similar terms that PC 2 enriched for, and positively enriched for transport associated and development terms. There is no enrichment for anterior specific function.

**Supplemental Fig 7.**
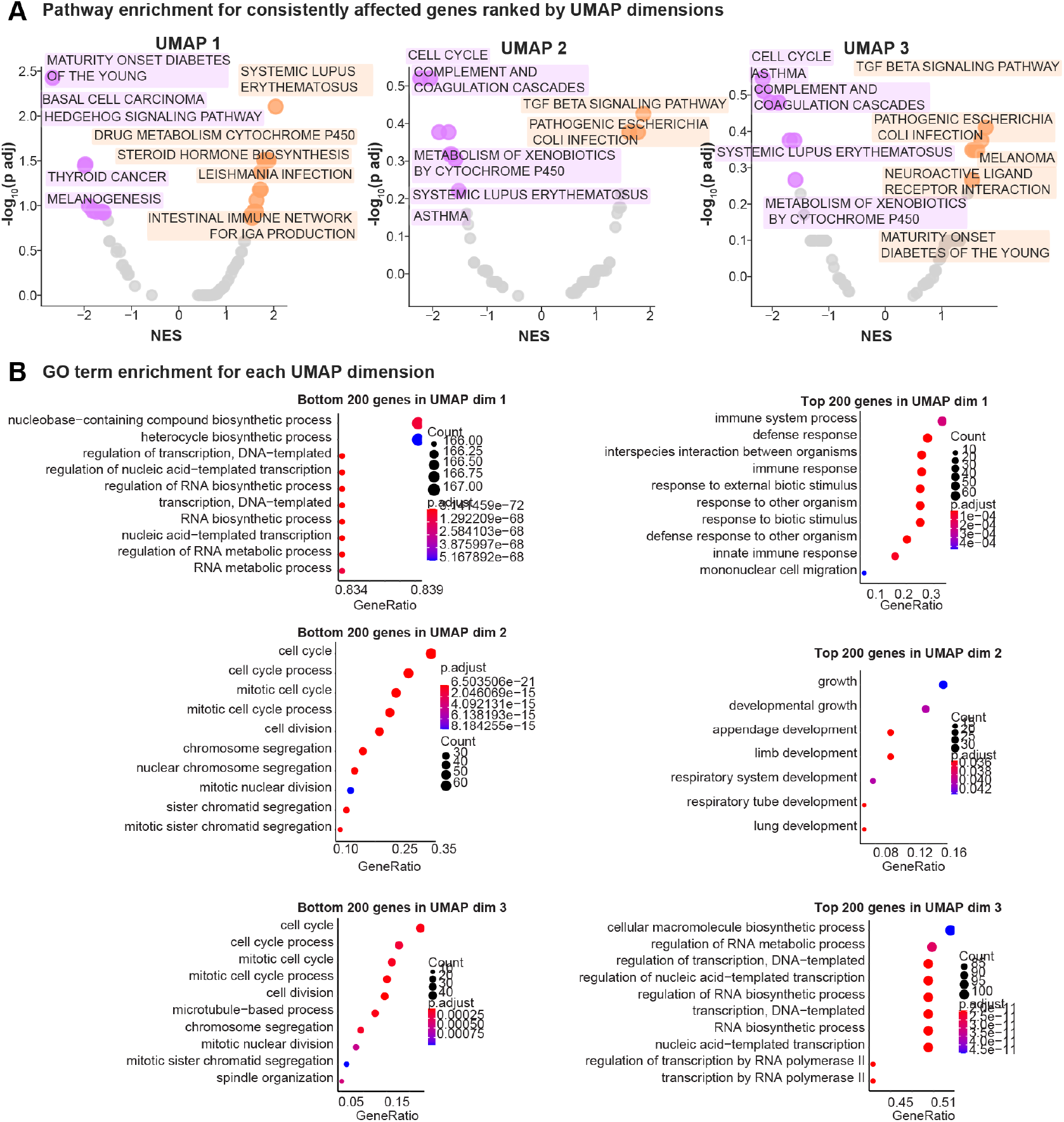
Enrichment for UMAP. (A) Genes were ranked by each UMAP dimension (up to 3) and to identify negatively and positively enriched pathways. UMAP 1 is negatively enriched for cancer associated pathways, and positively enriched for immune response. UMAP 2 is negatively enriched for a diverse range of pathways, including “cell cycle” and the immune associated “systemic lupus erythematosus”. UMAP 2 is only enriched for two pathways: “TGF beta signalling”, which is associated with development, and “pathogenic Escherichia coli infection”. The top five positive pathways in UMAP 3 covers similar pathways to UMAP 2 i.e., “cell cycle” and immune associated pathways. UMAP 3 is positively enriched for terms from UMAP 1 and UMAP 2. (B) The top and bottom 200 genes along each UMAP dimension were tested for enriched GO terms. UMAP 1 is positively enriched with RNA metabolism associated GO terms (contain the Hox genes), which does not overlap strongly with the enriched pathways. The top genes from UMAP 1 predominantly enrich in immune response terms and agree with pathways. The bottom genes from UMAP 2 are associated with cell cycle terms, while the opposing side of the dimension are development associated. In UMAP 3 both bottom and top terms overlap with UMAP 2. There is no enrichment for anterior specific function.

**Supplemental Fig 8.**
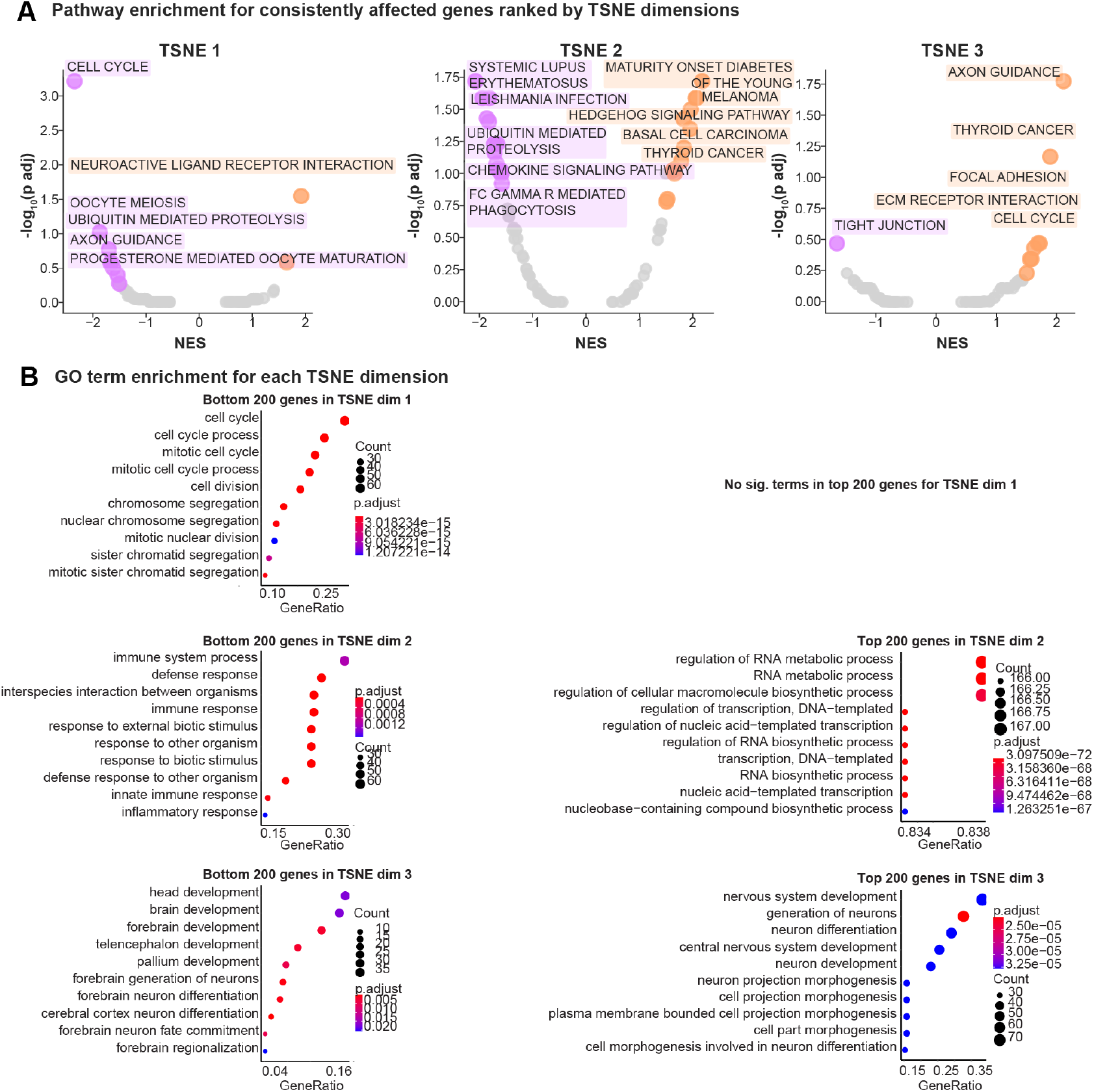
Enrichment for tSNE. (A) Genes were ranked by each tSNE dimension (up to 3) to identify negatively and positively enriched pathways. TSNE 1 is negatively enriched for cell cycle associated pathways and positively enriched for one pathway: “neuroactive ligand receptor interaction”. tSNE 2 is negatively enriched for immune response pathways and positively enriched for cancer associated pathways. TSNE 3 is negatively enriched for one pathway; “tight junction”, while positively enriched for diverse terms, including “cell cycle” and brain pathways. (B) The top and bottom 200 genes along each tSNE dimension were tested for enriched GO terms. TSNE 1 aligns with cell cycle associated terms, fitting with the enriched pathways along dimension 1, while not enriched for any terms in the top genes. In dimension 2, the bottom genes enrich immune response genes, and the top RNA metabolism associated GO terms (contain the Hox genes), all of which agree with enriched pathways. The terms for bottom genes in tSNE 3 overlap with forebrain development, while the terms for the top are more diverse, analogous to pathways, enriching for cell projection as well as development terms. None of the gene cohorts appear to directly encode for development terms.

**Supplemental Fig 9.**
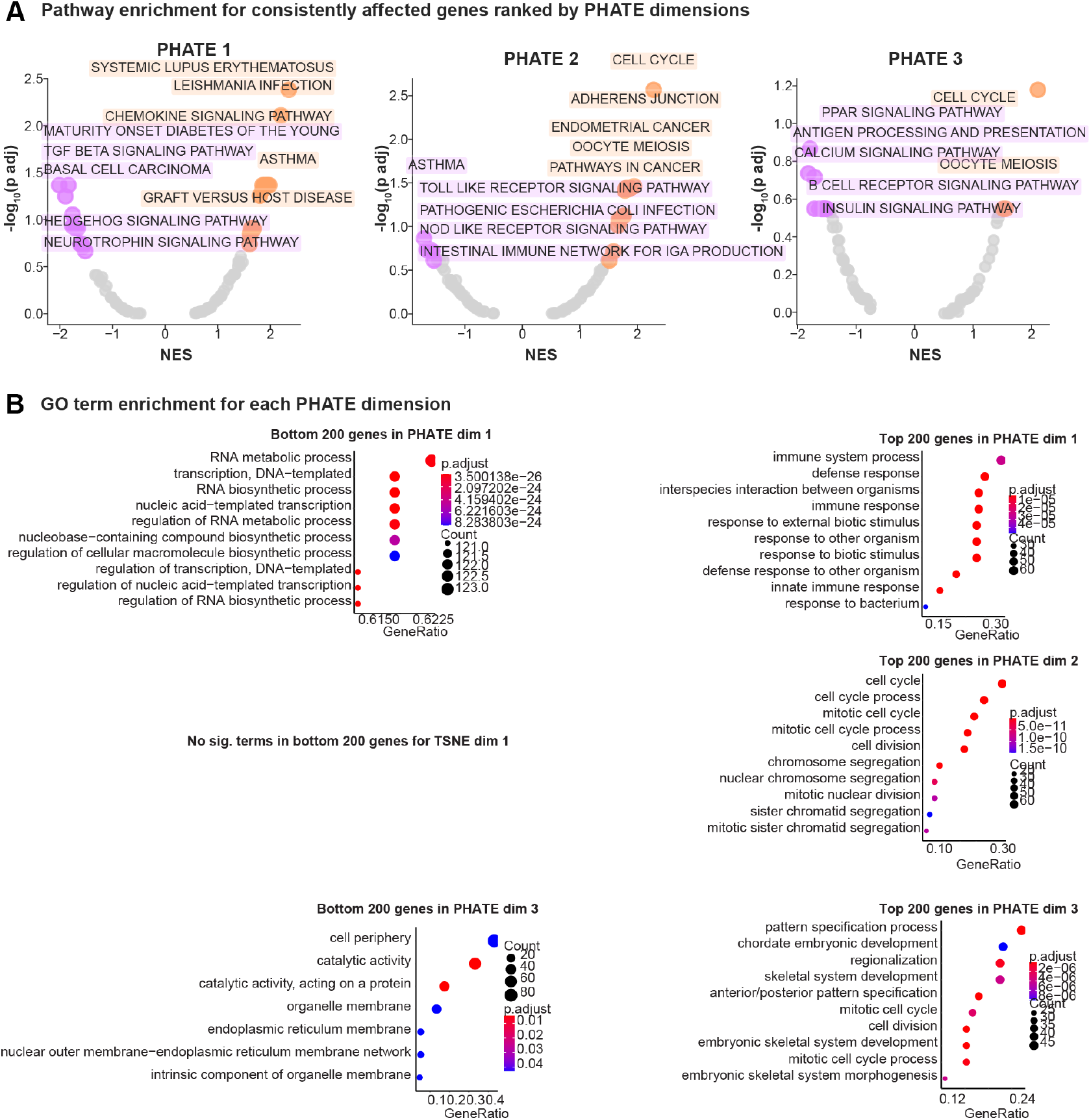
Enrichment for PHATE. (A) Genes were ranked by each PHATE dimension (up to 3) to identify negatively and positively enriched pathways. PHATE 1 is negatively enriched in cancer and brain associated pathways and positively enriched in immune associated pathways. PHATE 2 is negatively enriched in immune response pathways and positively enriched in cancer and cell cycle associated pathways. PHATE 3 is negatively enriched in signalling pathways and positively enriched in cell cycle pathways. (B) The top and bottom 200 genes along each PHATE dimension were tested for enriched GO terms. PHATE 1 bottom genes enrich for RNA metabolism associated GO terms (contain the Hox genes), the top genes of this dimension are enriched for immune response. In dimension 2 the top genes enrich for cell cycle terms similarly to the pathways in (A). The bottom genes from PHATE 3 overlap with cell periphery, and membrane terms, these terms are not enriched in any of the other methods as such would be a unique group. The top genes are associated with development, which does not overlap with the most significant pathways (cell cycle). There is no enrichment of the groups for anterior specific function.

**Supplemental Fig 10.**
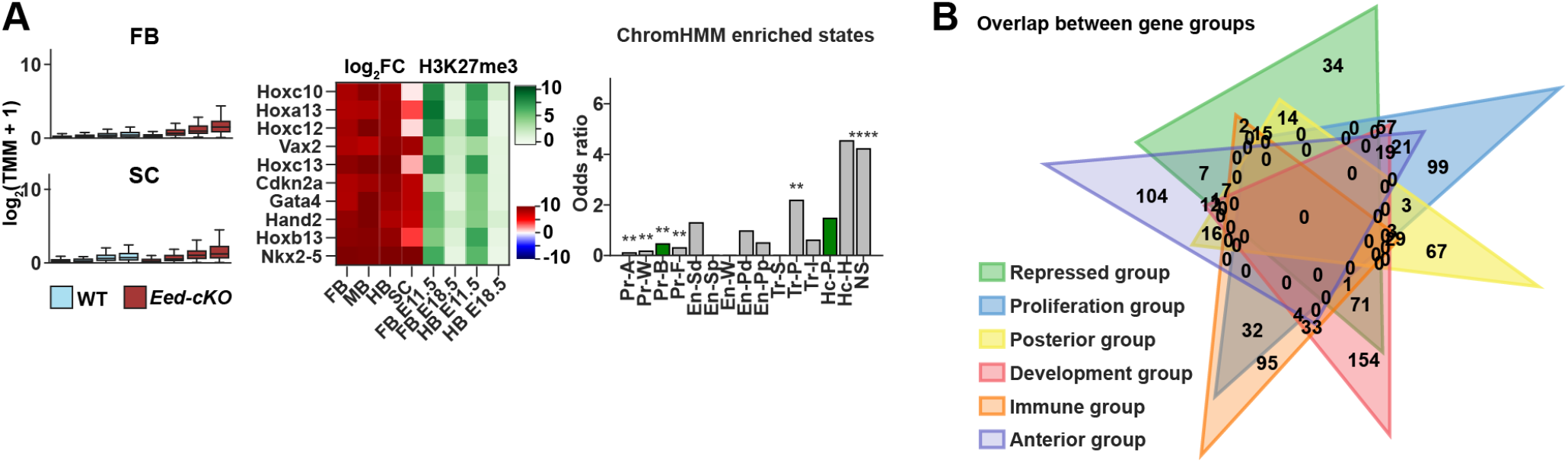
Results for repressed cohort. (A) Genes from the repressed gene cohort show overall low gene expression in both FB and SC. They are marked and enriched for the no-signal (NS) ChromHMM chromatin state. (B) Overlap between extreme groups of genes identified by the VAE shows that the repressed group strongly overlaps with the posterior group.

**Supplemental Fig 11.**
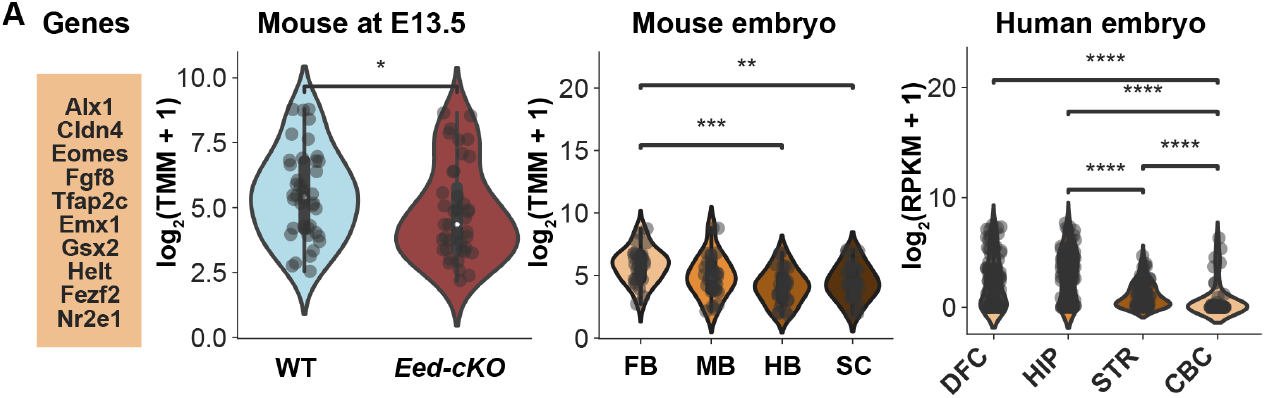
Evolutionary conservation into humans. (A) The top 10 genes from the anterior cluster showed conserved trends across human and mouse data.

